# *Plasmodium falciparum* Kelch13 and its artemisinin-resistant mutants assemble as hexamers in solution: a SAXS data driven shape restoration study

**DOI:** 10.1101/2021.02.07.430181

**Authors:** Nainy Goel, Kanika Dhiman, Nidhi Kalidas, Anwesha Mukhopadhyay, Ashish, Souvik Bhattacharjee

## Abstract

Artemisinin-resistant mutations in PfKelch13 identified worldwide are mostly confined to its BTB/POZ and KRP domains. To date, only two crystal structures of the BTB/POZ-KRP domains as tight dimers are available, which limits structure-based interpretations of its functionality. Our solution Small-Angle X-ray Scattering (SAXS) data driven shape restoration of larger length of protein brought forth that: i) PfKelch13 forms a stable hexamer in P6 symmetry, ii) interactions of the N-termini drive the hexameric assembly, and iii) the six KRP domains project independently in space, forming a cauldron-like architecture. While artemisinin-sensitive mutant A578S packed like the wild-type, hexameric assemblies of dominant artemisinin-resistant mutant proteins R539T and C580Y displayed detectable differences in spatial positioning of their BTB/POZ-KRP domains. Lastly, mapping of mutations known to enable artemisinin resistance explained that most mutations exist mainly in these domains because they are non-detrimental to assembly of mutant PfKelch13 and yet can alter the flux of downstream events essential for susceptibility to artemisinin.

## Introduction

Malaria is caused by the protozoan parasites of the genus *Plasmodium. Plasmodium falciparum* is the most virulent species in the African region, while *P. vivax* contributes equally to the malaria epidemiology in South-East Asia (SEA). In 2018, an estimated 228 million cases of malaria occurred globally with an estimated 405,000 deaths [1]. The parasite’s intraerythrocytic asexual cycle mediates all the symptoms and pathologies associated with acute infection as well as severe disease and artemisinin (or its derivatives) are frontline therapy for both the maladies [2, 3]. Artemisinins are potent drugs, used in combination with longer lasting antimalarials (Lumefantrine/Amodiaquine/Sulfadoxine/Piperaquine) as Artemisinin-based Combination Therapy (ACT). ACTs has significantly contributed towards reducing the global burden of malaria in the last two decades [4, 5]. However, the rapid emergence and spread of resistance to artemisinins threatens the worldwide malaria control and eradication strategies [6, 7]. Resistance to artemisinins is now widespread in the Greater Mekong subregion (GMS), including Cambodia, Yunnan province of China, Lao People’s Democratic Republic, Thailand, Vietnam, and Myanmar [8, 9]. Ironically, parasites in these countries also show increased resistance to the ACT partner drugs such as piperaquine and mefloquine, thus mitigating a higher rate of ACT failure [10, 11]. A potential risk of spreading of artemisinin-resistance from GMS to the Indian and African sub-continent, mimicking the earlier spread of chloroquine- and sulfadoxine/pyrimethamine-resistance, makes the control and eradication of this disease extremely challenging [12, 13].

Mutations in the *P. falciparum* PfKelch13 constitute the primary determinant of artemisinin resistance [14, 15]. These mutations alter several key pathways in the parasite’s intraerythrocytic cycle, including an overall reduction in the total ubiquitinated protein profiles and an increased expression of genes involved in unfolded protein response (UPR) pathways [16-18]. At the molecular level, artemisinin-resistant Cambodian isogenic strains with PfKelch13 mutations show a two-fold reduction in PfKelch13 abundance, a feature not exhibited by the two isogenic African strains [16, 19]. Artemisinin-resistant parasites also exhibit a constitutive basal level of PK4-mediated phosphorylation of *P. falciparum* eukaryotic initiation factor 2α (*Pf*eIF2α) in young rings but not in the young rings of artemisinin-sensitive Dd2 parasites [20]. Disruption of PfKelch13 function in the blood stages by conditional mislocalization using diCre-based gene deletion leads to a ring-stage arrest followed by slow transition into condensed parasite form, consistent with a similar arrest seen in the clinical artemisinin-resistant phenotypes [21, 22]. Recent reports also indicate that parasites with inactivated PfKelch13 or with artemisinin-resistant mutations display reduced hemoglobin uptake by endocytosis, thus diminishing the level of heme-induced activation of artemisinin to dihydroartemisinin (DHA) and thereby resulting in the induction of resistance [23-25]. Overexpression of mutant of wild-type PfKelch13 in resistant parasites was found to restore susceptibility thereby suggesting that artemisinin-resistant PfKelch13 mutations cause a loss-of-function. A proposed PfKelch13 substrate is the parasite phosphatidylinositol 3-kinase (PfPI3K), an enzyme converting phosphatidylinositol (PI) to phosphatidylinositiol 3-phosphate (PI3P). PfKelch13-facilitated ubiquitination and degradation of PfPI3K decreases the cellular PI3P level; and artemisinin-resistant PfKelch13 mutants display decreased ubiquitination and degradation of PfPI3K and an increased abundance of PI3P which may influence host cell remodeling and undermine artemisinin-induced proteopathy [16, 26]. These studies offer a good summary of the cellular response to artemisinin and its resistance; however, they fail to address a key question: What are the conformational changes in the overall PfKelch13 structure as it incorporates various mutations in an effort to mitigate this resistance?

PfKelch13 is encoded by *pf3d7_1343700* gene located in the chromosome 13 of the 3D7 strain. The PfKelch13 protein of 726 amino acids is comprised of a disordered and poorly conserved Apicomplexa-specific N-terminus region (amino acids 1-58) followed by three highly conserved and annotated domains, namely: a coiled-coil domain (CCD; amino acids 241-308); a Broad-complex, tramtrack and bric-à-brac; (BTB)/ the poxvirus and zinc-finger (POZ) domain (amino acids 350-452) and a C-terminal Kelch-repeat propeller (KRP) domain (amino acids 444–725) with six kelch motifs/blades (amino acids 444-490, 492-539, 540-574, 585-615, 633-680 and 682-725) (**Figure 1a**). Most of the PfKelch13 mutations detected in SEA are confined only to the KRP domains [14, 15]. However, worldwide polymorphisms in PfKelch13 spans the entire sequence [14, 15, 27, 28]. PfKelch13 C580Y and PfKelch13 R539T are the most dominant alleles in SEA while the most frequent PfKelch13 mutation in Africa is A578S, which has been subsequently validated as artemisinin-sensitive both *in vivo* and *in vitro* [8, 29]. Other nonsynonymous PfKelch13 mutations associated with delayed parasite clearance in the GMS are F446I, Y493H, I543T, P553L, R561H, P574L, A675V [30, 31]. In PfKelch13, the BTB and KRP domains display classical features of the multi-subunit RING-type E3 ligase complexes, thus suggesting that PfKelch13 functions as a substrate adaptor for E3 ubiquitin ligases. Ubiquitination of specific substrates is brought about by the recruitment of Cullin (which interact with the E3 ligases) via the BTB domain and substrate by the KRP. The E3 ligase activity then catalyzes the transfer of ubiquitin from the ubiquitin-conjugating enzyme (E2) to the protein substrate(s) and ultimately results in its proteasomal-mediated degradation [32]. Comparative structural and evolutionary analyses of the BTB domain of PfKelch13 clusters it with the BTBs of Potassium (K+) Channel Tetramerization Domain (KCTD) protein family with the highest similarity to KCTD17 [33]. Small-angle X-ray scattering (SAXS) studies and crystal structures of various KCTDs reveal a variety of stable oligomerization assemblies that allow efficient Cullin3 binding [34]. However, the oligomerization geometry of PfKelch13 in solution is largely unknown. The KRP domain of PfKelch13 exhibits a conserved, rigid, solvent-exposed shallow pocket that is similar to other KRP domain containing proteins, e.g., KEAP1, KLHL2, KLHL3 and KLHL12 [25].

**Figure 1.**
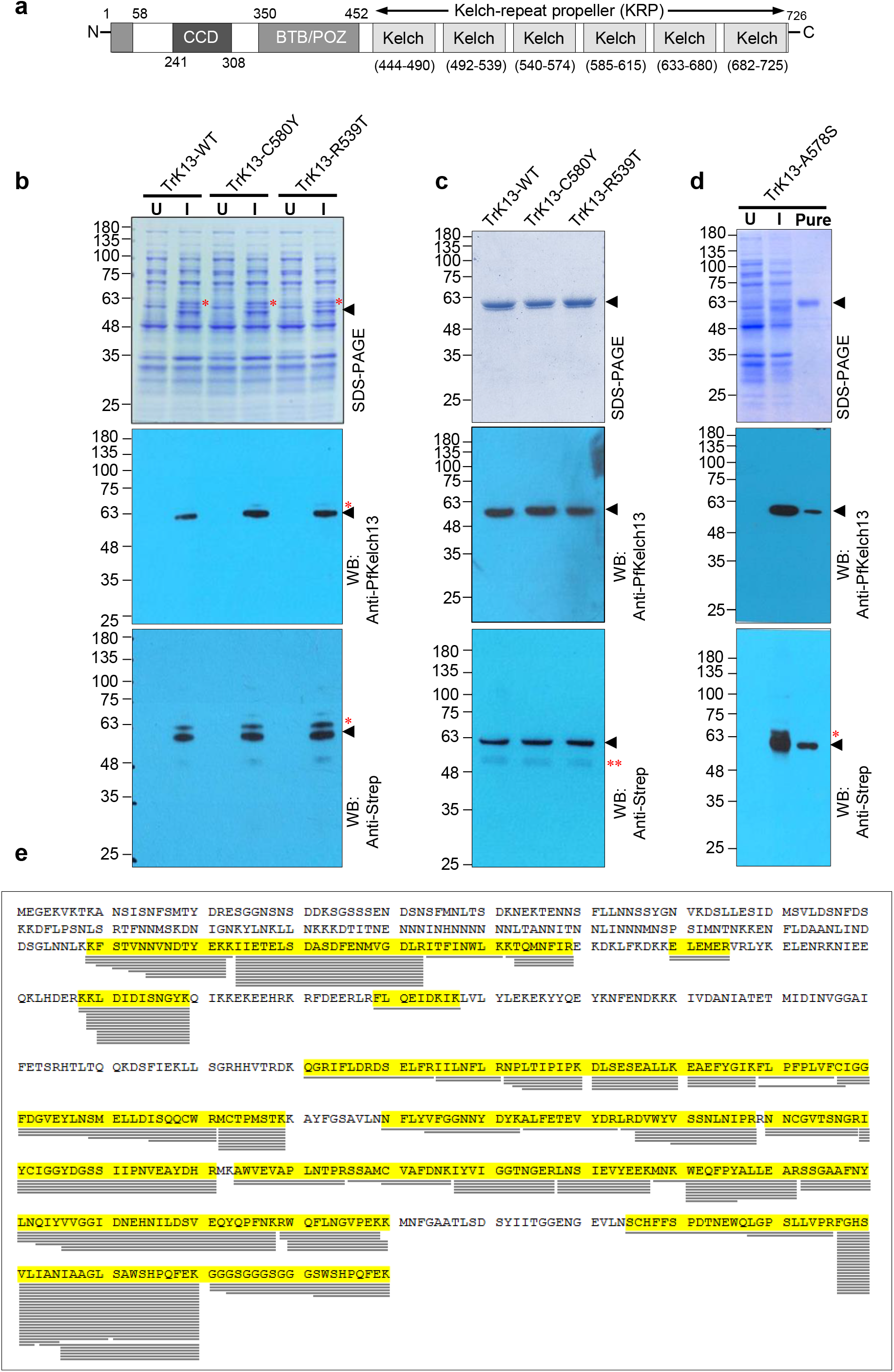
Expression and purification of recombinant TrK13-WT, TrK13-C580Y and TrK13-R539T. **a.** Schematic representation of PfKelch13 showing the Apicomplexa-specific N-terminus (1-58 amino acids), CCD (241-308 amino acids), BTB/POZ region (350-452) and the KRP domain (444-725 amino acids) with the six individual β-propeller/Kelch regions. **b**. SDS-PAGE (top panel) and western blots (middle and bottom panels) of *E. coli* cells showing the preferential expression of truncated TrK13-WT, TrK13-C580Y and TrK13-R539T at ∼61-kDa (arrowheads) on induction with ITPG (I), as compared to the respective uninduced control (U). Red asterisks indicate minor processed products of size >61-kDa. **c**. Analyses of purity and specificity of the affinity-purified TrK13-WT, TrK13-C580Y and TrK13-R539T by SDS-PAGE (top panel) and western blots (middle and bottom panels) using antibodies to PfKelch13 (middle panel) or Strep tag (bottom panel). Double red asterisks (bottom panel) indicate minor degraded products that are not recognized by PfKelch13 antibodies (middle panel). Molecular weight standards (in kDa) are as indicated. **d**. SDS-PAGE (top panel) and western blots (middle and bottom panels) of *E. coli* cells showing the preferential expression of truncated TrK13-A578S at ∼61-kDa (arrowheads) and recognition by anti-PfKelch13 (middle panel) or anti-Strep (bottom panel) antibodies only in samples induced with ITPG (I), as compared to the uninduced control (U). The purity and specificity of the affinity-purified TrK13-A578S by SDS-PAGE (top panel) and western blots (middle and bottom panels) are also shown. Red asterisks indicate minor processed products of size >61-kDa. **e**. LC-MS/MS analyses of TrK13-WT after trypsin digestion indicating the positions and lengths of detected peptides. The most N-terminus peptide detected is KFSTVNNVNDTYEK.

To date, only two crystal structures of PfKelch13 are available in RCSB Protein Data Bank, *i*.*e*., 4YY8, spanning amino acids 349-726 and 4ZGC, representing amino acids 338-726. While 4YY8 is predicted to be a monomer, 4ZGC is a disulfide-linked dimer [35]. Analyses of these solid-state structures as tight dimers has initiated interpretations that most PfKelch13 mutations are clustered in two specific regions, with the surface-exposed residues implicated in protein-protein interactions, and the buried residues influencing the overall PfKelch13 structure [27, 36]. However, both 4YY8 and 4ZGC lack a large segment of PfKelch13, thus limiting insights into the contributions of the N-terminus and the CCD in the overall structure and assembly. In this study, we expressed and purified recombinant wild-type PfKelch13 and three mutants using *E. coli* expression system. Two of these mutants (C580Y and R539T) represent dominant artemisinin-resistant mutations seen worldwide, while the third (A578S) is artemisinin-sensitive. Our SAXS data analyses and shape restorations allowed us to deduce the predominant solution shape of wild-type PfKelch13, the level of its association and how it is altered upon single point mutation.

## Results

### A truncated recombinant PfKelch13 is expressed and purified from *E. coli*

PfKelch13 is encoded by the intron-less *pf3d7_1343700* (*pfkelch13*) gene. The full-length *pfkelch13* was cloned into the bacterial expression plasmid pPR-IBA101 (IBA, Germany) under the control of T7 promoter and fused *in-frame* with sequences encoding for one-Strep affinity tag, a tandem repeat of 8 amino acids (WSHPQFEK) separated by glycine-rich linker, at the C-terminus. Cloning also engineered two alanine and a GL (glycine-leucine) ‘spacer’ between the coding end of PfKelch13 and the beginning of the one-Strep tag to ensure independent folding of PfKelch13 and the one-Strep tag. Surprisingly, the SDS-PAGE profile on IPTG induction revealed a preferentially induced band at ∼61 kDa rather than the expected mass of ∼87-kDa for the full length strep-tagged PfKelch13 (**Figure 1b**, top). Standardizations of various conditions by altering either the time of induction, lowering the induction temperature (to as low as 16°C), or the media composition (Luria broth *versus* terrific broth) did not significantly alter the expression profile and the induced band was consistently seen at ∼61-kDa (data not shown). Western blot analyses using commercial antibodies to either one-Strep-tag (IBA, Germany) or custom-generated anti-PfKelch13 (Genscript, USA) confirmed the ∼61-kDa protein as strep-tagged PfKelch13, *albeit* a N-terminally truncated product (**Figure 1b**, middle and bottom). We thus designated the ∼61-kDa **Tr**uncated and strep-tagged recombinant Pf**K**elch**13** as **TrK13-WT** (wild-type) during this entire study.

Transformed *E. coli* AI cells expressing recombinant PfKelch13 with R539T, C580Y (the dominant mutations seen in clinically artemisinin-resistant parasites) and A578S (artemisinin-sensitive mutation, widely seen in Africa) mutations also showed specific induction/accumulation of ∼61-kDa proteins (instead of their respective ∼87-kDa full-length counterparts) similar to the TrK13-WT in SDS-PAGE and western blots (**Figures 1b, d**). Accordingly, these were named as TrK13-C580Y, TrK13-R539T and TrK13-A578S, respectively in this study. The induced TrK13-WT, TrK13-C580Y, TrK13-R539T and TrK13-A578S were affinity-purified using StrepTactin XT resin, dialyzed against PBS, pH 7.4 and concentrated using Amicon concentrators. SDS-PAGE and western blot analyses using antibodies to PfKelch13 confirmed the purity and specificity of these recombinants (**Figure 1c**). In addition, western blot using anti-Strep antibodies also showed the presence of a smaller truncated product (<5% densitometry-based calculated intensity as compared to TrK13-WT, TrK13-C580Y and TrK13-R539T; data not shown) at ∼53-kDa in all (**Figure 1c**, double asterisks in the bottom). These might represent non-specific processed fragments of TrK13-WT, TrK13-C580Y and TrK13-R539T, respectively from amino acids 311-726 (plus one-Strep tag) that are devoid of the anti-PfKelch13 epitope and with theoretical predicted molecular weight of 50980-Da (*www.expasy.org*). It is pertinent to mention here that since the research manuscripts on crystal structures 4YY8 and 4ZGC are not available to date, and the details on the protein expressions, post-purification status and constructs used for crystal set-up are unknown. Particularly, it is unknown if these protein versions were also susceptible to degradation, as seen by us.

Independent digestion of TrK13-WT by trypsin or AspN followed by LC-MS/MS revealed the most N-terminus peptides as KFSTVNNVNDTYEK(K) or KFSTVNNVN, respectively, where the first lysine residue corresponds to the 188^th^ amino acid position of the full length PfKelch13 (**Figure 1e** and **Supplementary Figure 1**). LC-MS/MS data also suggested that the processing at the N-terminus in *E. coli* was entirely non-specific, since other heterogenous peptides such as ^189^FSTVNNVNDTYEK, ^190^STVNNVNDTYEK, ^192^VNNVNDTYEK and ^194^NDTYEK peptides were also detected at the N-terminus (**Figure 1d** and **Supplementary Figure 1**). The differences in molecular masses of these Trk13-WT variants were too small to be resolved by conventional SDS-PAGE. The purified TrK13-WT was estimated to be between 534-538 amino acids in length (excluding the 34 amino acids spacer and one-Strep tag sequences). The predicted molecular mass of the TrK13-WT is ∼66-kDa and we speculated that the aberrant mobility was likely due to the unusual amino acid composition, as previously described for many recombinant *Plasmodium* proteins [37]. The basis for the specific accumulation of TrK13-WT (rather than full-length PfKelch13-strep) in bacterial cells was unclear.

To address the truncation of TrK13-WT, we bioinformatically predicted of protease cleavage sites in full-length PfKelch13 by PeptideCutter (*www.expasy.org*). The output revealed the presence of low-specificity Chymotrypsin/LysN/Pepsin/Proteinase K and LysC/LysN/Trypsin cleavage sites at 187^th^ and 188^th^ amino acid positions, respectively (**Supplementary Table 1**). However, these proteases are reportedly absent in *E. coli* AI cells, which are derived from *E. coli* BL21 strain and are also deemed Lon and OmpT protease deficient. We also investigated if the N-terminus of full length PfKelch13 was inherently disordered and stimulated a non-specific enzymatic digestion in the bacterial cells. *In silico* analyses of the PfKelch13 sequence by both the Protein Disorder Prediction System (*PrDOS; http://prdos.gc.jp*) [38] and IUPred2A (*https://iupred2a.elte.hu*) indicated a greater propensity of disorder at the N-terminus of PfKelch13 as compared to the downstream sequences (**Figures 2a, b** and **Supplementary Table 1**). Specifically, PrDOS predicted semi-continuous disordered stretches from amino acids 1-65, 125-143, 154-181, 266-306. The predicted disordered region spanning amino acids 266-306 was located within the coiled-coil domain (241-308) with a defined secondary structure and thus was not considered. However, the semi-continuous disordered stretches at the N-terminus till amino acid 181 could induce a bacterial-driven auto-catalytic event that resulted in a truncated TrK13-WT. Similarly, IUPred2A interfaces two different servers, namely IUPred2 and ANCHOR2, to identify disordered protein regions, respectively [39]. For the full-length PfKelch13 (with C-terminus one Strep tag; 759 amino acids), IUPred2A output indicated a disordered stretch at the N-terminus 1-174 amino acids and globular conformation for amino acids 175-738 (including the one-Strep tag) (**Figure 2c**).

**Figure 2.**
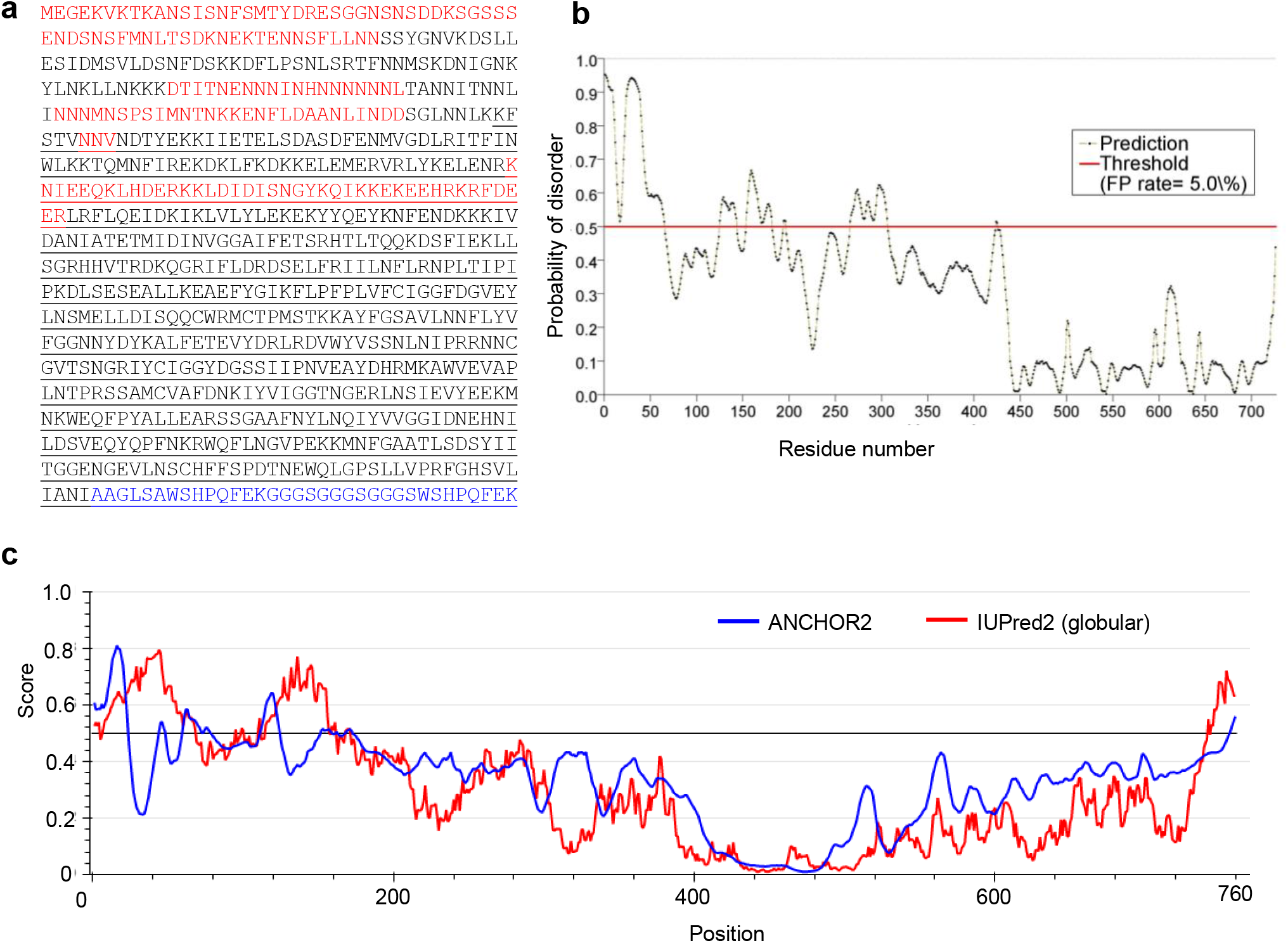
Higher disorder propensity at the N-terminus of PfKelch13. **a-b.** Amino acid sequence of the full-length PfKelch13-WT-strep showing the disordered stretches (a, red text), as predicted by PrDOS (*http://prdos.hgc.jp*) and its graphical representation **b**. The sequence of TrK13-WT is underlined. The spacer and the one-STrEP tag are shown in blue text. **c**. IUPred2A (*https://iupred2a.elte.hu*) analyses of PfKelch13-WT-strep indicating residues with higher disorder scores are mostly confined to the N-terminus of PfKelch13-WT-strep and precedes the N-terminus of recombinant TrK13-WT, which starts from amino acid 188 onwards.

### SAXS data analyses from solutions of TrK13-WT, TrK13-R539T, TrK13-C580Y and TrK13-A578S reveal hexameric assembly

To date, two crystals structures of C-terminal portion of PfKelch13 covering the BTB and the KRP are available (PDB IDs: 4YY8 and 4ZGC) (unpublished) [35]. The refined chains in the asymmetric units implied that these fragments of protein were parallel dimers with tight packing of their BTB and KRP domains. Interestingly, the chains in the PDB deposition ID 4ZGC were interlinked by an artificially engineered S-S linkage which obviously influenced the observed assembly. Since most of the mutations that render resistance to artemisinin were located in BTB-KRP of PfKelch13, these dimers have been used to perceive functional role of the corresponding residue replacement in the PfKelch13 structure [27, 33, 36]. Interestingly, none of the known resistance rendering mutants have been yet analyzed experimentally by any structural biology method. Thus, to understand PfKelch13 structurally, we were able to express and purify TrK13-WT, TrK13-R539T, TrK13-C580Y and TrK13-A578S as described in the methods. For all the affinity purified proteins, gel filtration experiments with Superdex200 10/300 GL column showed as a single peak eluted just after the void volume. SDS-PAGE of the peak fraction(s) resolved as a single band at ∼61-kDa in each case (data not shown). These observations supported that TrK13 proteins show some degree of association of 300-kDa or higher in molecular mass. Additionally, reinjection of portions of purified peak led to the same profile in gel filtration experiments, *albeit* with lower intensity. In this study, at the foremost we acquired solution SAXS data at 10°C with three concentrations of freshly purified TrK13-WT *i*.*e*., 1.4, 0.8 and 0.25 mg/ml and the data collected from protein samples from three different batches of expression-to-purification yielded comparative results (**Supplementary Figure 2**). Double log plots confirmed lack of any aggregation or inter-particulate effect in all the samples. Scaled plots of the concentration dependent datasets further suggested any absence of influence of protein concentrations in the range studied, and the peak profile of the dimensionless Kratky plot confirmed globular nature of protein molecules in solution (**Supplementary Figure 2**). Linear region of the Guinier plot and residuals implied mono-disperse nature of scattering species. To gain structural insights in resistance-inducing mutations along with TrK13-WT, SAXS data were also collected on purified TrK13-R539T and TrK13-C580Y under identical conditions (**Figure 3**). In addition, data was acquired from a null mutant TrK13-A578S. All protein samples were in the concentration range of 1.1 to 1.5 mg/ml. The main plot confirmed: i) lack of aggregation or inter-particulate effect in all samples, and ii) overall profile of all, *i*.*e*., TrK13-WT and mutants, were similar thus supporting comparable large shape parameters for all the proteins (**Figure 3**). The latter outcome was also supported by the similar profile and position of the peaks in their Kratky plot (**Figure 3a**, inset). Further, dimensionless Kratky plots from all the proteins showed that the peak was below qR_g_ value of 1.73 for TrK13-WT and TrK13-A578S, and around 1.73 for TrK13-R539T and TrK13-C580Y, thereby suggesting similarities in the overall scattering shape of TrK13-WT and TrK13-A578S but differed in TrK13-R539T and TrK13-C580Y (**Figure 3b)**. Linear fits to the Guinier plots of all datasets supported monodisperse nature of protein molecules in all cases. Guinier analysis presuming globular and rod-shaped profiles for TrK13-WT, TrK13-R539T, TrK13-C580Y and TrK13-A578S provided R_g_ and R_c_ values of 5.77, 6.77, 6.42 and 5.9 nm, and 3.78, 3.22, 3.9 and 3.75 nm, respectively. Using the relationship between R_g_ and R_c_, persistence length, L of the molecules were estimated to be 15.1, 20.6, 17.7 and 15.8 nm, respectively. This processing of the log q data indicated that R539T and C580Y mutations induce increase in the length of the protein molecules as compared to TrK13-WT.

**Figure 3.**
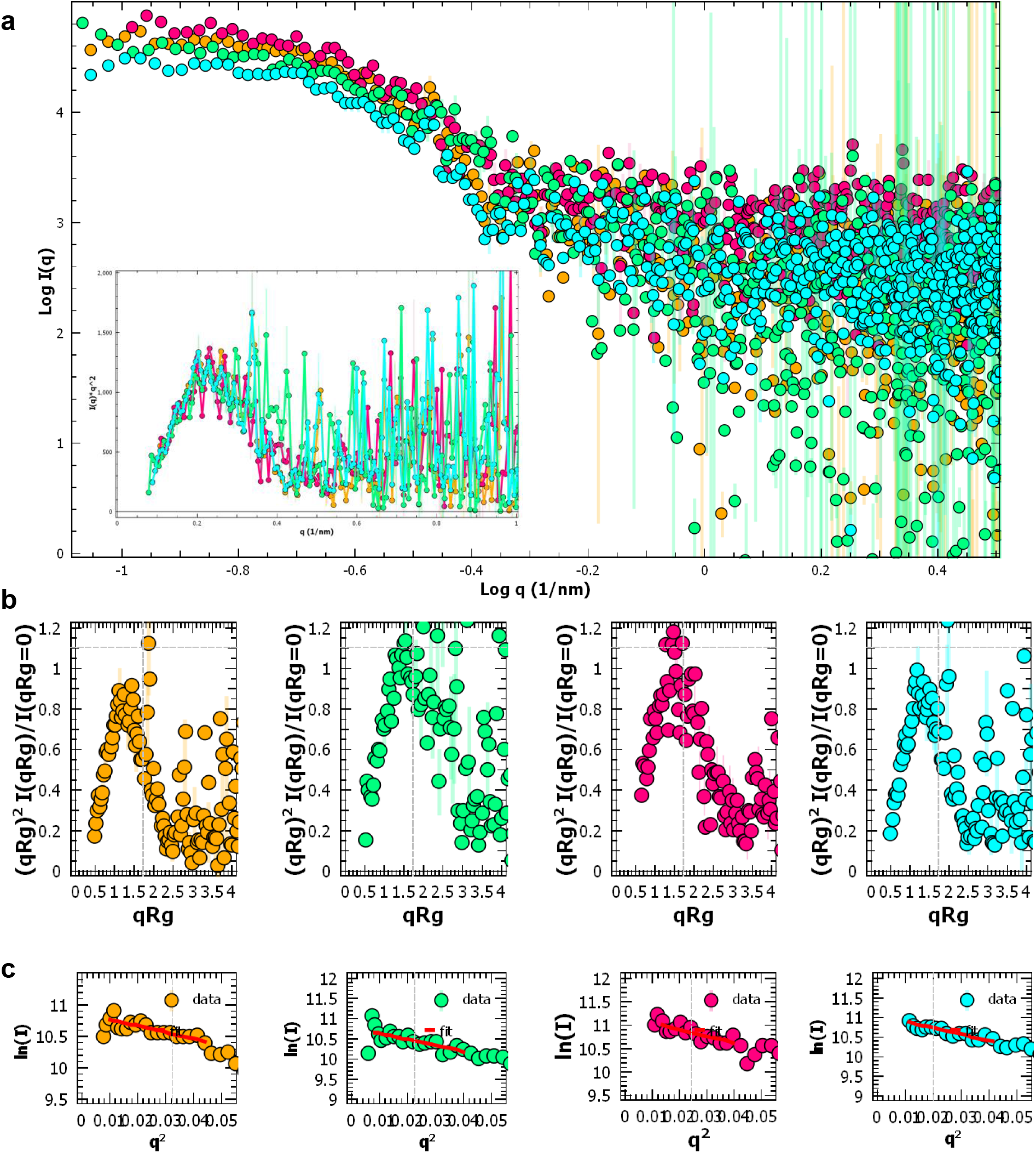
SAXS data from TrK13-WT, TrK13-R539T TrK13-C580Y and TrK13-A578S. **a.** Double Log plot of the SAXS datasets acquired from samples of TrK13-WT (orange), TrK13-R539T (green), TrK13-C580Y (magenta) and TrK13-A578S (cyan) are shown. Inset shows the Kratky plots of the same datasets. **b**. Dimensionless Kratky plot of the datasets is shown at same scale. **c**. Linear fits (red line) to the Guinier analysis of respective datasets presuming globular shape profile are presented.

Using a wider range of q, the distance distribution profiles of interatomic vectors inside the SAXS profiles, P(r) were estimated (**Figure 4**) for all proteins. The estimated curves depicted shoulder and peak like profile at approximately 5.5 and 10 nm for TrK13-WT and TrK13-A578S, respectively. The main peak was at lower r value for TrK13-R539T and TrK13-C580Y. The analysis also provided D_max_ and R_g_ values of TrK13-WT, TrK13-R539T, TrK13-C580Y and TrK13-A578S as 17.1, 22.1, 18.4, 16.8 nm, and 6.6 ± 0.09, 6.9 ± 0.11, 6.8 ± 0.09 and 6.7 ± 0.07 nm, respectively. The trends were similar to estimations from Guinier region, thereby suggesting that resistance inducing R539T and C580Y mutations were leading to increment in dimensions of molecules; however, non-effective mutant A578S folded similar to the WT. The analysis also provided I_0_ values from the samples which, in correlation with values obtained from standard protein samples and estimated concentration of these samples, provided a molecular mass of TrK13-proteins in the range of 359-385 kDa. Additionally, Bayesian calculations of molecular mass from SAXS datasets indicated mass of scattering species in solution to be about 320-345 kDa. Since theoretical mass of all the TrK13 proteins at single chain level was ∼61 kDa, both the approaches supported a hexameric association level of proteins in solution for TrK13-WT and mutants. These results correlated with the higher mass peak seen in the gel filtration studies. So, we performed additional size exclusion chromatography experiments using SHODEX Protein KW-800 GFC column attached to HPLC system [40]. Individually, all the TrK13 proteins eluted as single peak in between Apoferritin (443-kDa) and β-amylase (200-kDa) suggesting a mass of about 340-350-kDa for the TrK13 proteins (data not shown). To rule out any low temperature induced hydrophobic collapse or influence on association status, we also acquired SAXS datasets at different temperatures ranging from 10°C to 60°C (**Supplementary Figure 3**). Stacking of the SAXS datasets acquired at intervals of 10°C confirmed that the hexameric assemblies of TrK13-WT and mutants did not induce any heating induced aggregation or any significant changes in the solution shape. Kratky plots of the datasets further confirmed retention of globular shape for all proteins in the temperature range studied. Overall, these analyses confirmed that the hexameric assemblies intrinsic to PfKelch13 were stable and independent of the tested mutations.

**Figure 4.**
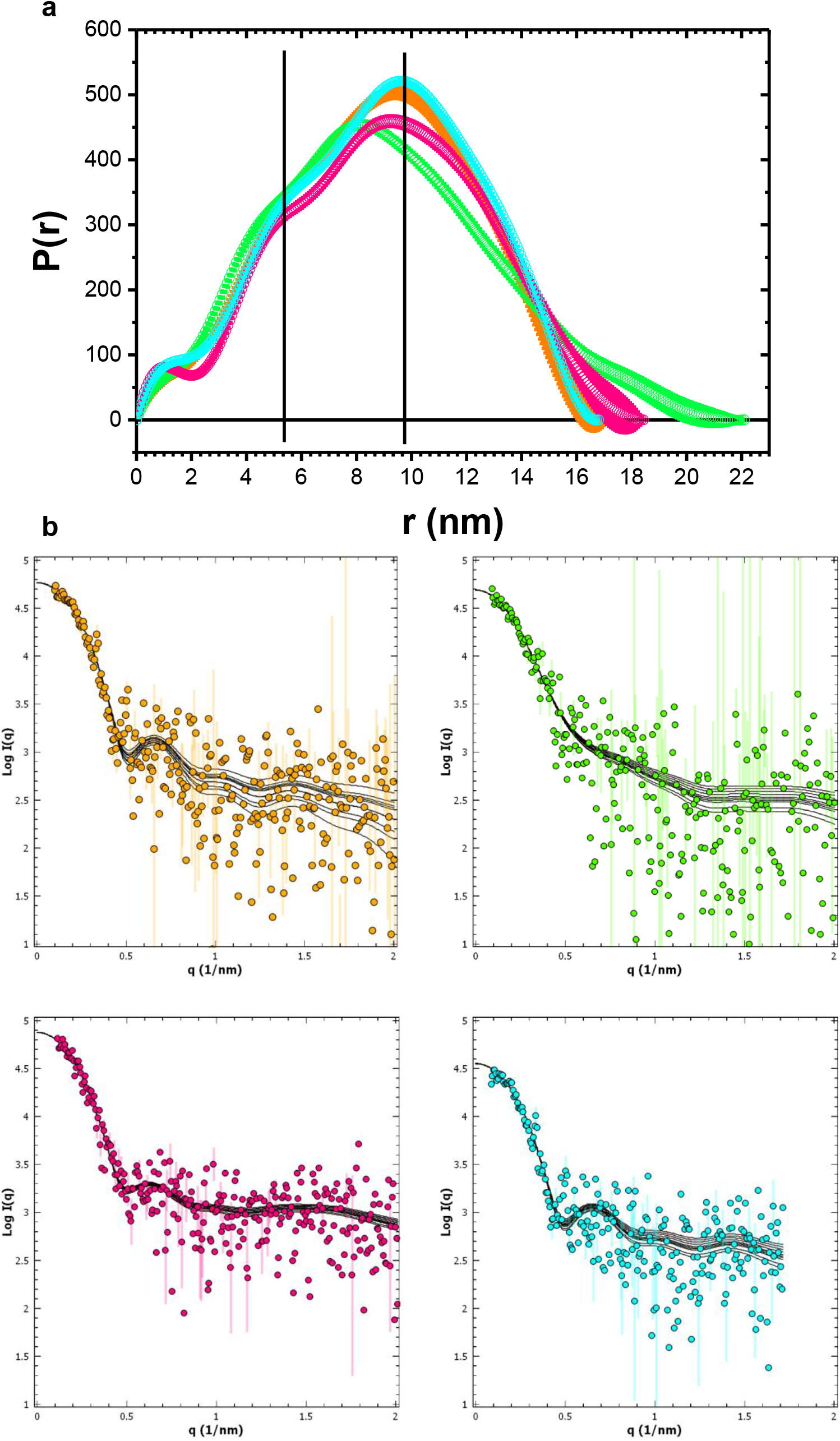
Distance distribution curves and profiles of computed models. **a.** The computed distance distribution profiles, P(r) of the interatomic vectors inside the shape of the TrK13-WT (orange), TrK13-R539T (green), TrK13-C580Y (magenta) and TrK13-A578S (cyan) from their SAXS datasets are plotted. **b**. Comparison between the theoretical SAXS profiles (black lines) of the ten models solved for each protein with the respective experimental data for TrK13-WT (top left), TrK13-R539T (top right), TrK13-C580Y (bottom left) and TrAK3-A578S (bottom right).

### Shape Restoration of the TrK13-WT and mutants

Using the respective SAXS data profile and estimated P(r) curves, we generated ten uniform density models and averaged (**Figure 4a**). Calculated resolution of the solved models varied in range of 5.3 to 6.4 nm, and the estimated NSD values between models were 0.534, 0.621, 0.78 and 0.672 for TrK13-WT, TrK13-R539T, TrK13-C580Y and TrK13-A578S, respectively. Black lines in the plots (**Figure 4b)** depict theoretical SAXS profiles of the ten individual models solved for each protein assembly. Different views of the molecular maps of the average model generated for TrK13-WT and mutants are shown in **Figure 5**. The different panels in the first, third and fifth rows depict the cauldron-shaped association of the proteins in solution, as solved in P6 symmetry, and suggests that all the protein assemblies are similarly associated on one face (fifth row) and have a hollow feature on the other side (first row). The side-view (third row) of these proteins clearly indicates increased length of the association for TrK13-R539T and TrK13-C580Y (demarcated by reference parallel dotted lines) as compared to the TrK13-WT and TrK13-A578S. The automated overlays with respect to TrK13-WT (shown in rows second, fourth and sixth) help in realizing that the main differences between these proteins are in the shape of the domains bordering the opening as a function of single point mutations in the proteins (best visualized in the first and second rows).

**Figure 5.**
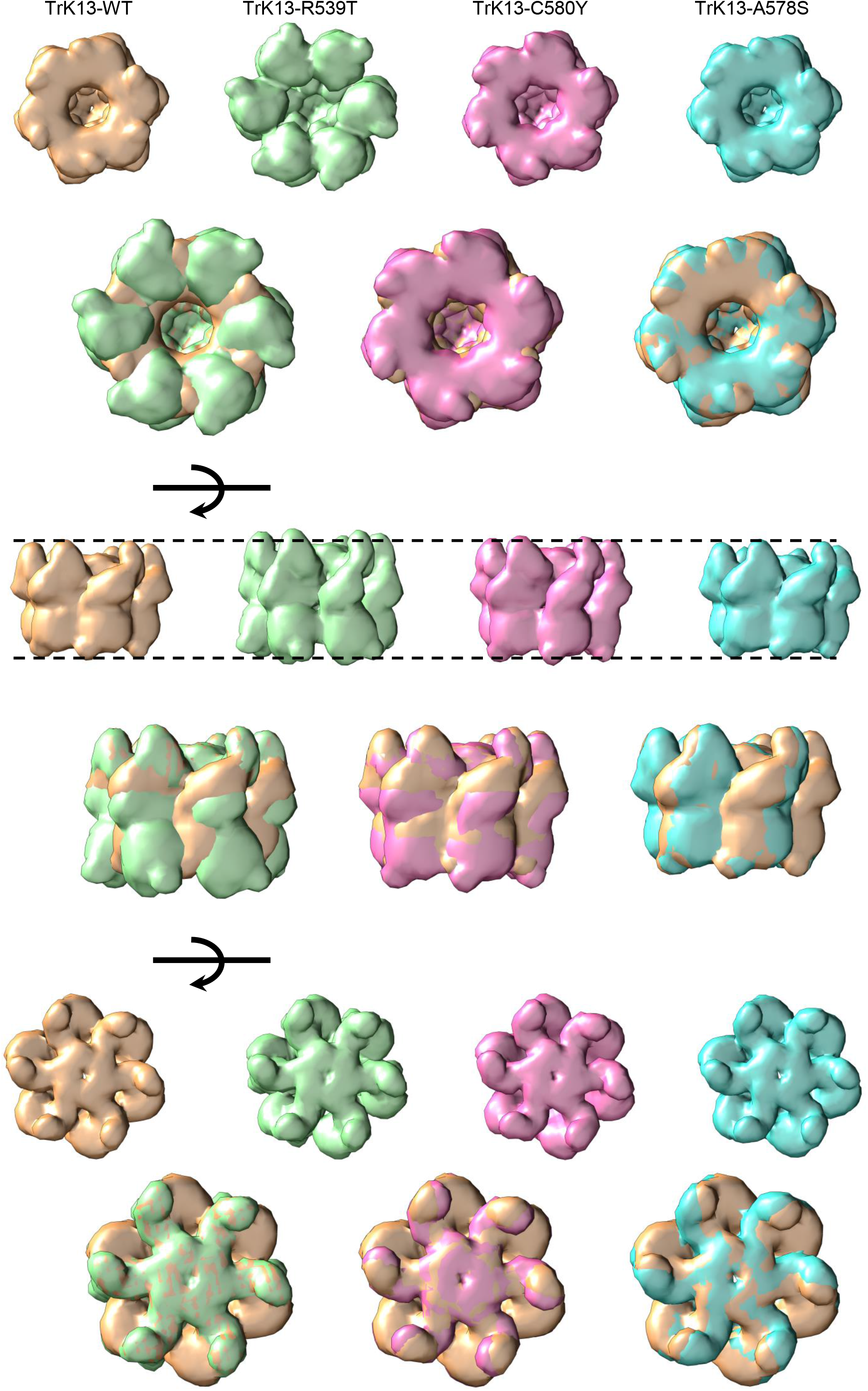
Molecular shapes of the averaged model of ten solutions for each protein. First row shows one view of the shape envelopes of all protein which highlights the hollow cauldron feature retained in TrK13-WT (orange), TrK13-R539T (green), TrK13-C580Y (magenta) and TrK13-A578S (cyan). **Second row** shows the inertial axes alignment of the shapes solved for mutant protein versus wild type. **Third and fourth, and fifth and sixth rows** display rotated views of shapes individual proteins and their respective overlaps with TrK13-WT. The black arrows indicate the orthogonal rotation used to generate the view presented.

In the absence of any reference crystal structure of full-length PfKelch13 (4YY8/4ZGC incorporate only the BTB-KRP regions), we generated a 3D atomic model of TrK13-WT using the Phyre2 server (*www.sbg.bio.ic.ac.uk/phyre2*) based on remote homology and template-based assembly simulations to the existing crystal structures in the PDB [41] (**Supplementary Figure 4a**). As described, to distill residue definition model of TrK13 proteins, we divided the primary structure of TrK13-WT (and the mutants) into six parts and allowed each fragment to translate and rotate in the 3D space freely within the constraints of their end-to-end connectivity as per the sequence and SAXS data based on the profile of their respective model (**Supplementary Figure 4b, c**). The dummy residue based SAXS data profile constrained models were manually dissected in 1/6^th^ part to represent chain volume residing in one chain of TrK13-WT or mutant. Mass estimation of the representative models of chain using CRYSOL program was about 57-59 kDa for the proteins. The theoretical shape profile of the dummy residue model of single chain of each protein was then used to reposition the six fragments of TrK13 protein to generate a solution where the calculated SAXS profile of spatial positioning of fragments agreed with previous profile using SASREF [42]. These segments were stitched together by template driven modeling using primary structure of the protein and SWISS MODELER server (*https://swissmodel.expasy.org/*) [43]. The net end-result was an energy-optimized and continuously ligated structural model of single chain of TrK13-WT or mutant proteins. For each protein, six copies of this structural model of single chain were positioned in space using experimental SAXS dataset as a reference to generate hexameric models of the proteins using SASREF program. The automated placement of the residue scale hexamer model inside the envelope of the model solved using dummy residues confirmed that the blobs or domains at the mouth of the opening of the hexameric assemblies were formed by the KRP domains while the N-terminal helices appeared to seal the other side of the cauldron (**Figure 6**, rows 1-4). Views in the fifth row portray the residue scale models of the four hexamers in ribbon format, and the clippings of same shown in the sixth row highlight that mutations in the KRP domains induce changes detectably in the KRP region of the assembly, while the N-terminal helices remain unchanged in the Trk13-WT and mutants. The packing of individual chains in their hexameric assemblies and the secondary structural element in that chain are shown (**Figure 7a**). When superimposed, the individual chains matched the chain in the hexameric assembly deduced by experimental SAXS data. **Figure 7b** shows the comparisons of the single chains for individual mutant *versus* TrK13-WT. It is pertinent to mention that no structural alignment has been done between the chains. The arrows 1 and 2 highlight the helix and β-sheet rich BTB and KRP domains in the structural models. These comparisons implied that although mutations in TrK13-R539T and TrK13-C580Y were in the KRP domains, their net perturbations were detected throughout the BTB domain but not across the linker regions leading to the N-terminal of the assembly. In contrast, these changes were minimal for TrK13-A578S which fared like theTrK13-WT at the BTB-KRP regions. For the R539T and C580Y mutations, the KRP domains seemed to extend away in the long axis of the protein structure and oriented slightly at an angle. Thus, from this aspect of modeling we concluded that the artemisinin resistant mutations induced detectable conformational changes in their respective BTB-KRP domains. These perturbations disseminated throughout their molecular assemblies and were subsequently ‘absorbed’ by the flexible linker upstream of the BTB region that resulted in their gradual decay as they approached the disordered N-terminus. As a result, the net effect was barely detectable at the extreme N-terminal end of the molecules. Since the N-terminal portions are the driving force for association, the susceptible and resistance mutants remained hexameric despite point mutations in the lower part of protein.

**Figure 6.**
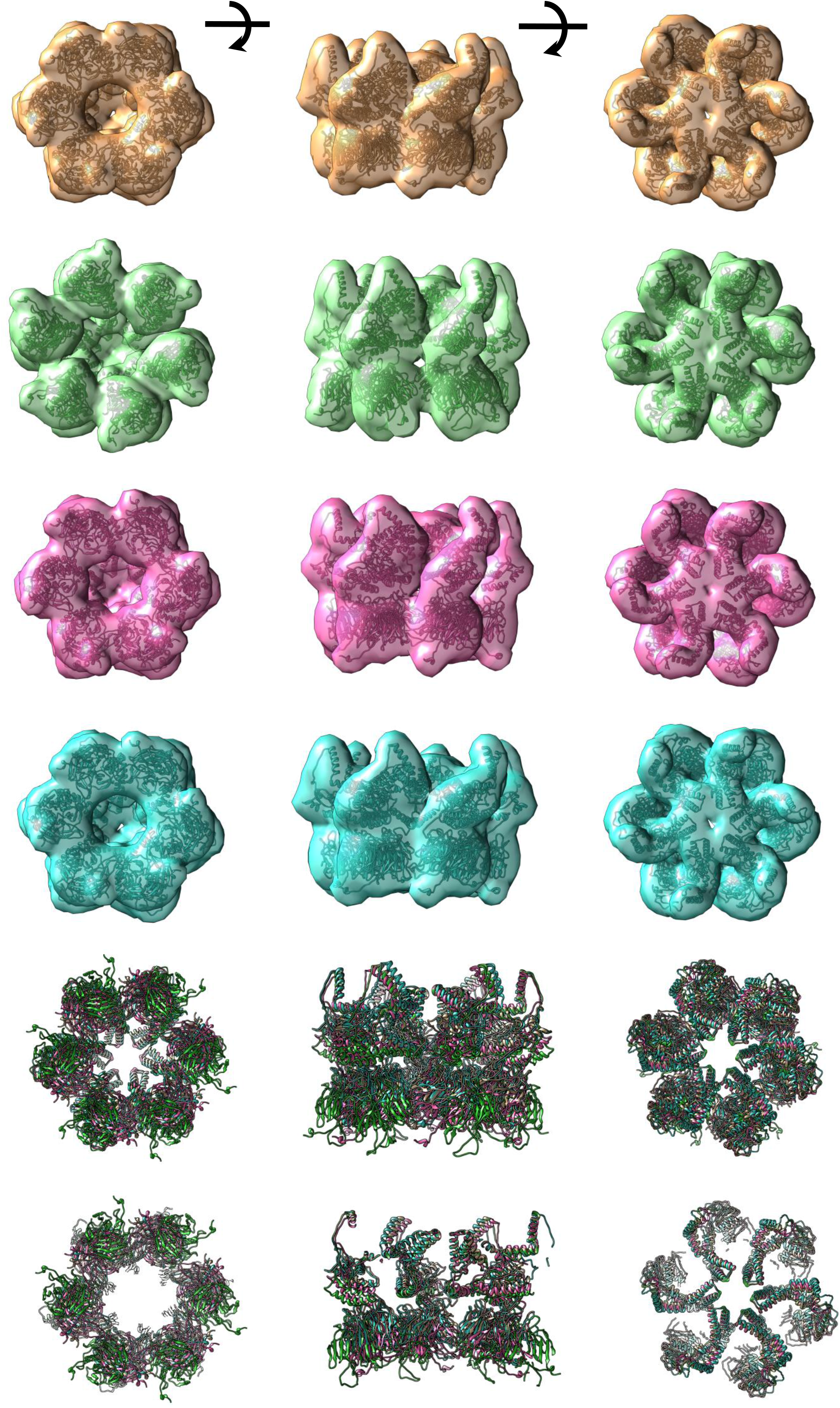
Comparison of molecular shapes from SAXS data with their residue detail models. In **top four rows**, envelopes shown in Figure 5 are superimposed with residue detail models solved using segments of domains of proteins oriented in space using SAXS data (ribbons). Color codes are same as Figure 5. **Fifth and sixth rows** provide a look at ribbon models solved as placed inside SAXS data-based envelopes. Left panels have the KRP domains facing the plane of view, mid panels are rotated side views, and right panels represent the N-terminal side of the assembly facing the plane of view. In **sixth row**, for left image, clipping was done from the back of page to fade out N-terminal part and show only BTB-KRP domains of all proteins in their assemblies. In mid image, clipping was done from front to show transverse section of associations, and in right image, clipping was again done from back to fade out BTB-KRP domains to show how N-terminal helices interact in assemblies.

**Figure 7.**
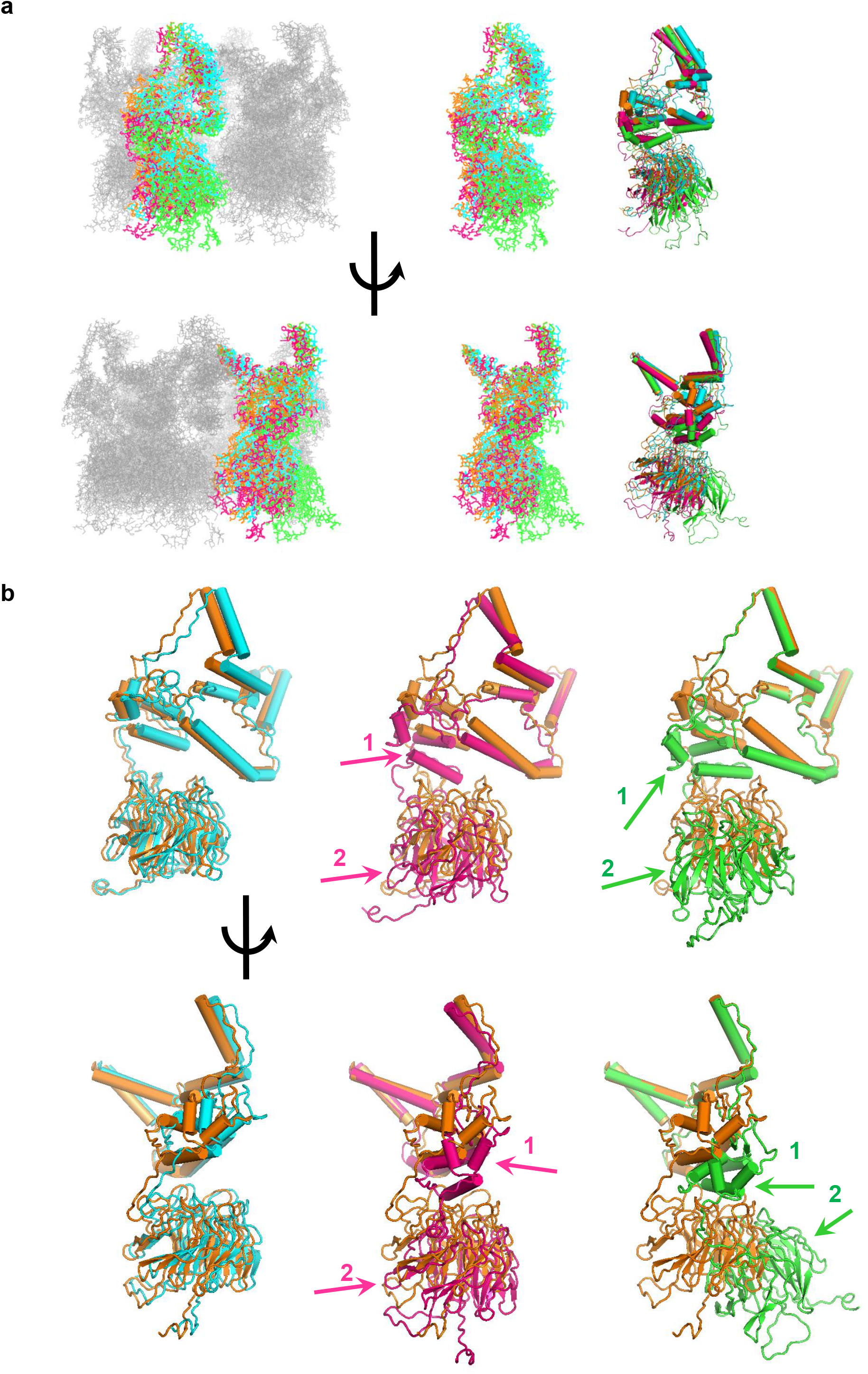
Understanding structural changes between individual chains in the protein assemblies. a. Left images are two in plane rotated views of one chain of all proteins in their SAXS data based hexameric assembly have been highlighted [TrK13-WT (orange), TrK13-R539T (green), TrK13-C580Y (magenta) and TrK13-A578S (cyan)]. Other chains have been shown as grey lines. In right panels, single chain has been shown in line mode and highlighting secondary structural content. **b**. Overlap of single chains in the models have been shown to highlight local changes between wild type and mutant protein as per modeling. Arrows 1 and 2 point to BTB and propeller domains in the overlap seeking attention to repositioning of these domains in the mutant protein chains particularly in TrK13-R539T (green) and TrK13-C580Y (magenta) versus TrK13-WT (orange).

## Mapping of the landmark artemisinin-sensitive and -resistant mutations in the PfKelch13 hexamer

Most of the non-synonymous mutations in PfKelch13 reported worldwide are confined to the KRP domain [14, 28, 29, 44, 45]. We were interested in visualizing if the relative positioning of these mutations in PfKelch13 were altered in the changed conformations of the hexameric TrK13-R539T and TrK13-C580Y, as compared to TrK13-WT. Since there was no single comprehensive report assigning a value to the efficacy of resistance for different mutations in PfKelch13, we referenced a recent meta-analysis report that compared the standardized estimates of the PC_1/2_ with PfKelch13 propeller region genotypes and assigned xPC_1/2_ values (where x corresponds to the probability that PC_1/2_>x is less than 0.05) as a cutoff for infections with ‘slow-clearing’ parasites [30]. The parasite clearance half-life (PC_1/2_) measures the slope of the log-linear component of the clearance curve for parasites with reduced susceptibility to artemisinins on day 3 following ACT and has been established as the best *in vivo* assay to validate artemisinin susceptibility [46]. We thus chose xPC_1/2_ values as a standard to group the different PfKelch13 mutations. Accordingly, we assigned the parasites with mutations in PfKelch13 with xPC_1/2_ in the range of <1.39, 1.4-1.69, 1.7-1.99 and ≥2 as sensitive (0), low-resistance (+), medium-resistance (++) and high-resistance (+++), respectively (**Supplementary Table 2**). The previously validated artemisinin resistant PfKelch13 mutations associated with slow parasite clearance (C580Y, R539T, Y493H, I543T, R561H and N458Y) were denoted +++ and the candidate PfKelch13 mutations (P441L, G449A, G538V, P553L, V568G, P574L and A675V) were designated ++. Parasites with significant differences in the PC_1/2_ values from different sites and studies (F446I, E252Q and A481V) were assigned as +. Other mutations, like A578S, K189T and K189N were not associated with artemisinin-resistance, and hence, were left unassigned [30]. These residues were highlighted across all the four hexameric assemblies (**Figure 8)**. Surprisingly, though the propeller domains were significantly altered in TrK13-R539T and TrK13-C580Y, we observed that the resistance-inducing residues (+++ and ++) retained their original positions at the outer periphery of the KRP assemblies similar to the TrK13-WT and TrK13-A578S. Clearly the R539T and C580Y mutations led to repositioning of KRP domains that caused the assembly to differ from one side, but the functionally key residues were not grossly mis-or re-oriented. Overall, these observations implied that the dominant artemisinin resistant C580Y and R539T mutations may instead alter a basic structural feature in PfKelch13, and this possibly rewires subset of its downstream functions that ensures survival of these parasites in the duration of ACT regimen or *in vitro* artemisinin treatment.

**Figure 8.**
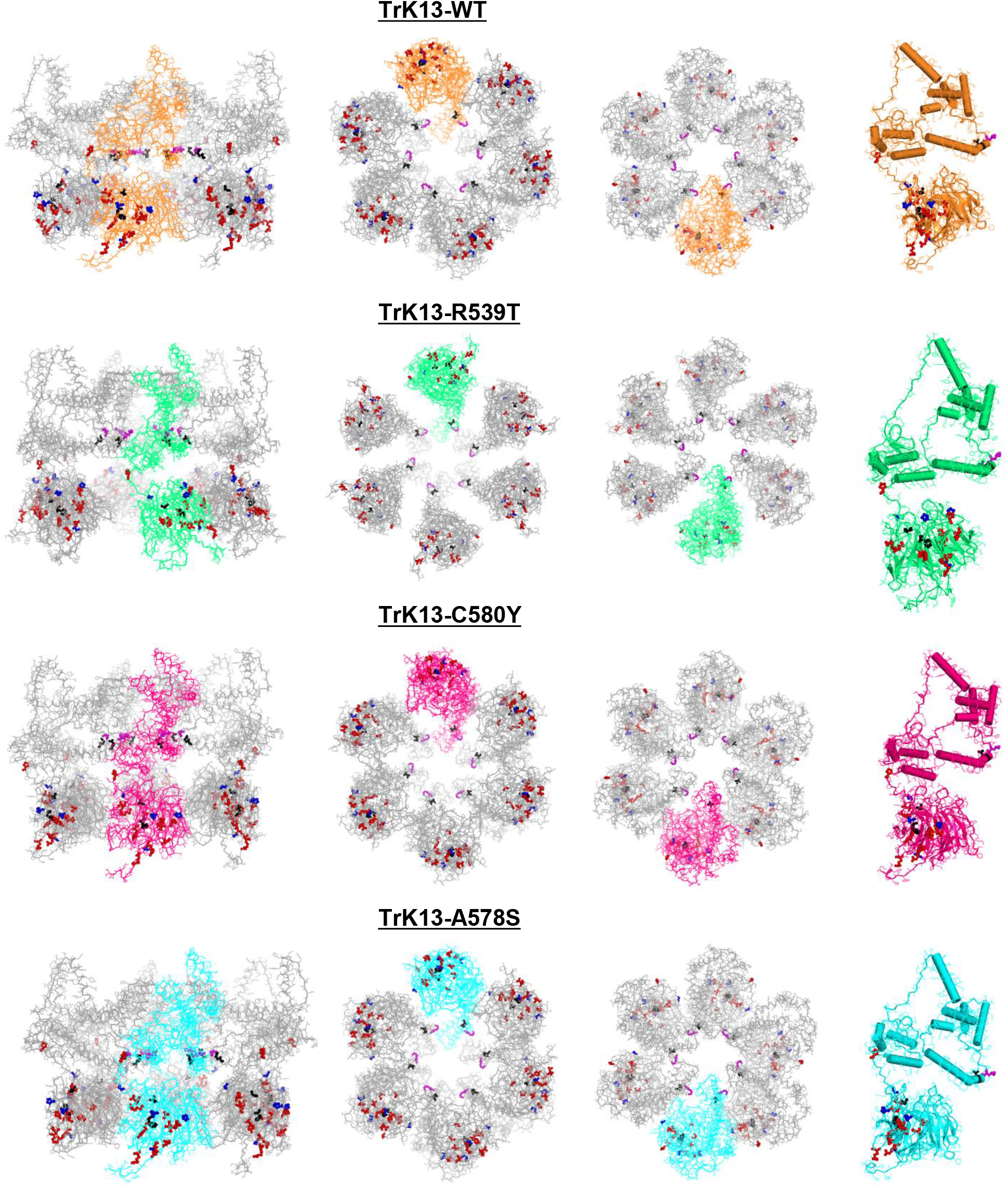
Mapping of spatial location of Kelch13 residues known to render artemisinin resistance in the hexameric assembly. Top to bottom lanes display rotated views of modeled assemblies of TrK13-WT, TrK13-R539T, TrK13-C580Y and TrK13-A578S. Individual chains are colored as in previous figures. Panels in first or leftmost column show the side view of the assemblies; second column panels project propeller domains to readers; third column has images where N-terminal portions are towards reader, and right column has image of single chain in secondary structure mode. Residues key to function are listed in Supplementary Table S2 and are shown as spheres. Relative sensitivity order has been colored as: +++ red; ++ blue; + black and 0 magenta.

## Discussion

Non-synonymous mutations in *P. falciparum* PfKelch13 are associated with resistance to artemisinins. Invariably, all the structural studies of PfKelch13 invoke either of the two available (yet unpublished) crystal structures in the PDB: 4YY8 (amino acids 349-726) and 4ZGC (amino acids 338-726), each with individual chain in the asymmetric unit packing tightly as a parallel dimer [35]. Both 4YY8 and 4ZGC have included only the BTB-POZ (amino acids 350-452) and the KRP (amino acids 444-726) domains, conveniently excluding other functionally relevant regions like the N-terminus and the CCD. However, lessons from prior studies have emphasized that evolutionary constraints generally prohibit random tethering of domains in any multi-domain protein; rather individual domains in such proteins bear an intrinsic property to modulate the structural features of neighboring domains *in trans* [47, 48]. We thus believed that the dimeric association seen in 4YY8 may actually be a conformational collapse induced by the absence of the N-terminus and CCD regions located upstream of the BTB or could have been induced by the lattice packing effects during crystallization. Not limited by need for diffraction quality crystal, the solution SAXS offered us a practical platform to investigate the conformational features of PfKelch13 wild-type (TrK13-WT) and its mutants in solution and obtain reliable insight into their shape properties.

Since the first discovery linking PfKelch13 to artemisinin resistance by Ariey and coworkers [14] and till date, to the best of our knowledge we are unaware of any published report on the recombinant expression of PfKelch13 using *E. coli* or other eukaryotic expression systems. In this study, we successfully expressed and purified the recombinant wild-type PfKelch13 (TrK13-WT), the two most-prevalent artemisinin-resistant mutants R539T and C580Y (TrK13-R539T and TrK13-C580Y, respectively) and a widely occurring yet artemisinin-sensitive mutant A578S (TrK13-A578S) using *E. coli* expression system. We observed preferential accumulations of truncated recombinants, with lengths spanning between amino acids 189-726 of the full-length PfKelch13, with 4-5 amino acid variations at the N-terminus, and independent of the presence of C580Y, R539T or A578S mutations (**Figures 1** and **Supplementary Figure 1**). We speculated that the disordered N-terminus might have prompted a non-specific enzymatic digestion in *E. coli* inducing such 4-5 amino acid variations [49]. This was consistent with our *in-silico* analyses using PrDOS and IUPred2A predictions whereby PfKelch13 sequence exhibited a greater propensity for disorder at the N-terminus, more specifically spanning the amino acids between 154-181 located just upstream of truncation site (**Figure 2**). The experiments in this study could not confirm activity by any inherent bacterial proteases, since the use of bacterial protease inhibitor cocktail did not affect the expression profiles (data not shown). Nonetheless, the purified TrK13-WT and mutants (TrK13-C580Y, TrK13-R539T and TrK13-A578S) retained most of the essential elements of full-length PfKelch13, including the truncated N-terminus-CCD-BTB-KRP to serve as a reliable template for studying the shape-function parameters and offer plausible correlation between the effect of each mutation on its overall structural conformation or assembly. In addition, recombinants used in our study featured marked improvement in PfKelch13 sequence, with ∼74.10% coverage in TrK13-WT and mutants, as compared to only ∼54% in 4YY8/4ZGC.

The SAXS profiles were similar across different batches or concentrations of purified proteins which ruled out their influence on any inter-particulate effect (**Figure 3** and **Supplementary Figure 2**). Surprisingly, Guinier estimations and Bayesian calculations for the interatomic distance distribution in conjunction with the relative protein concentrations of TrK13-WT and mutants, supported an intrinsic hexameric association in solution across TrK13-WT and mutants. These were not artifacts of expression or purification since they stably retained their association states and were inert to temperature-induced hydrophobic collapse or association (**Figure 4** and **Supplementary Figure 3**). Our independently generated uniform-density models ‘molded/shaped’ the hexamers into a cauldron-like architecture with each hexameric unit of TrK13-WT and mutants forming a tight association at the N-termini. Interestingly, our results showed that the KRPs at the C-termini were loosely organized and featured a hollow channel (**Figure 5** and **Figure 6**). These hexameric assemblies in TrK13-WT and mutants were consistent with the oligomerization states of the BTB domains of KCTD protein family where the PfKelch13-BTB domain has been reported to cluster structurally [33, 34].

In *P. falciparum*-infected erythrocytes, analytical microscopy data has previously localized PfKelch13 across diverse intra-parasitic destinations, including vesicular structures at or near the ER and plasma-membrane associated cytostomes [16, 21, 24]. PfKelch13 has also been reported to associate with mitochondrial proteins and the resident organelle post DHA (the active metabolite of all artemisinins) treatment and unaffected by the presence of R539T mutation [25]. In addition, biochemical fractionation experiments have indicated peripheral association of PfKelch13 with membranous structures and stereological analyses of cryo-immunoelectron microscopic sections localized PfKelch13 to both proximal and distal ER-tubules and vesicles [16]. Similar hexameric assemblies of proteins associated within the frameworks of vesicular trafficking across different systems are also not *hitherto* unreported. The N-ethylmaleimide-sensitive factor (NSF), involved in the disassembly of the soluble NSF attachment proteins receptor (SNARE) proteins; Synaptophysin I, an archetypal member of the MARVEL (MAL and related proteins for vesicle trafficking and membrane link)-domain family involved in fusion and recycling of synaptic vesicles and ERGIC-53, a member of L-type lectins exhibiting different intracellular distributions and dynamics in the ER-Golgi system of secretory pathway in eukaryotes are all known to display homo-hexameric assemblies [50-52]. The human heat-shock protein 90 (Hsp90) has been shown by TEM tomography to self-assemble into oligomers in the cytosol of mammalian cells upon heat shock or divalent cations treatment and may have a pivotal role in protecting cells from thermal damages by its chaperone function [53]. Interestingly, a recent study further reported volumetric resolution of N-terminally GFP-tagged PfKelch13 puncta at the surface of the parasite by 3D structured illumination microscopy (3D-SIM). The GFP-PfKelch13 structure revealed a doughnut-shaped architecture with a peak-to-peak intensity diameter of 165 ± 30 nm and reminiscent of cytostomal structures that are involved in uptake of host cell hemoglobin for the delivery to the parasite digestive vacuole [23]. We believe that the previously reported dimeric assembly of 4YY8 and 4ZGC do not justify the doughnut geometry effectively, instead the cauldron-like PfKelch13 hexamer identified in this study provides a reasonably fitting skeleton within the dimensions of the PfKelch13 doughnut visualized in 3D-SIM and may be consistent with cytostomal ring patterns [54]. We also observed increased longitudinal dimensions of the dominant artemisinin-resistant Trk13-R539T and TrK13-C580Y hexamers as compared to TrK13-WT and artemisinin-sensitive TrK13-A578S mutant (**Figure 5**). In this study, the conformational orientations of KRPs and the change in dimensions of the hollow channel were the main discriminating feature between the TrK13-WT and mutants. These changes may have resonated as an overall lengthening of the cauldron handle (represented by the CCD-BTBs) and ‘spreading-out’ of the cauldron diameter (constituted by the KRPs) in TrK13-R539T and TrK13-C580Y, a result also evident from our SAXS data. We also speculate that these changes might have affected the constitution of cytostomal ring, altering its dimensions and likely decreased the efficiency of hemoglobin uptake in parasites harboring such mutations. Such an effect probably impaired hemoglobin catabolism and heme-mediated activation of artemisinins and delayed in the onset of artemisinin activation, which might have translated to prolonged persistence of parasites with C580Y and R539T mutations in PfKelch13 during the presence of DHA. In fact, mislocalization of PfKelch13 in early ring-stage parasites has been previously reported to reduce hemoglobin-like peptides following exposure to DHA, essentially mimicking features of PfKelch13 mutant isolates [23]. Our interpretations also corroborate well with a recent quantitative colocalization microscopic study wherein PfKelch13-WT and PfKelch13-R539T mutant showed slightly increased and decreased Pearson’s Correlation Coefficient (PCC; considered as an indicator of association) values, respectively with three different markers of vesicular trafficking (Rab5A, 5B and 5C) immediately following DHA treatment and thereby suggesting decreased level of endocytosis in the R539T mutant following DHA exposure [25].

In this study, we manually fragmented the rigid Phyre2-generated TrK13-WT model into six distinct domains while taking careful considerations in maintaining their individual integrity (**Supplementary Figure 4**). This allowed us the ability to freely rotate each domain independently in the 3D space and stitch together with energy optimization to generate a single chain of TrK13-WT and replicate to six copies to position into the experimental SAXS dataset. The same method, when replicated across the mutants (TrK13-R539T, TrK13-C580Y and TrK13-A578S), translated the SAXS-based model profile in agreement with the theoretical shape profile of the dummy residue model. The net result clearly indicated that a single point mutation in the KRP induces profound changes only in the mouth of the cauldron, while the N-terminus helices constituting the cauldron base remain largely unaffected (**Figure 6 and 7**). We believe that this spatial association between the N-termini of different chains holding the hexamer together is critical for the integrity of PfKelch13 and may expand to its stability. This might have been imposed further by the evolutionary constraints that allows a greater propensity of disorder at the N-terminus of Pfelch13 (outputs from *in silico* PrDOS and IUPred2A predictions) to accommodate large variations in the KRP or BTB/POZ domains without affecting the structural integrity of PfKelch13 hexamer. Any mutation therein that compromises this hexameric assembly may gradually lead to its disintegration and misfolding of PfKelch13 and, eventually its proteasomal degradation. Moreover, *pfkelch13* gene has been established as essential for intraerythrocytic asexual growth of both *P. falciparum* and *P. berghei* parasites and conditional mislocalization of PfKelch13 or diCre-based gene deletion leads to an arrest at the early intraerythrocytic ring stage followed by slow transition to condensed parasite forms [21, 55, 56]. Immunoprecipitation experiments using PfKelch13-specific monoclonal antibodies has also recently been shown to enrich for multiple components of the 19S regulatory subunit of the 26S proteasome; thus, supporting the potential role of PfKelch13 as an adaptor for the delivery of polyubiquitinated proteins for proteasomal degradation [25, 57]. Alternatively, the possibility of self-targeting of disassembled or misfolded PfKelch13 for proteasomal degradation following ubiquitination cannot be ruled out.

Surprisingly, when we highlighted the PfKelch13 mutations implicated for the various degrees of artemisinin resistance across all the four hexamers, we observed that despite the KRP domains being significantly altered in TrK13-R539T and TrK13-C580Y, the relevant residues (and other functionally important mutants) retained their original position at the outer periphery of the propeller assembly as in TrK13-WT and TrK13-A578S (**Figure 8)**. It is noteworthy to state that mutations at the N-terminus of PfKelch13 occur only in sporadic instances with few known clinical examples worldwide. Commonly seen mutations include the non-synonymous K189T, K189N and synonymous K189K from Africa; double mutation H136N-C580Y from a single parasite clone in China, single case of I205T, E252Q from Western Thailand and Myanmar and 3 cases of R255K [30]. Among them, only E252Q had a smaller effect on parasite clearance with xPC_1/2_ value of 1.5, essentially classifying it as low-resistant (+) in our study. Epidemiologically, the lower resistant-conferring E252Q mutation has been prevalent in northwest Thailand-Myanmar border before the advent of the extremely resistant (+++) C580Y phenotype [30]. In our hexameric model of TrK13-WT, the K189, I205, E252 and R255 mutations are all localized at the inner interface of the TrK13-WT hexamer (H136 could not be highlighted in our study because of the truncated TrK13-WT) near the vicinity of the extreme N-terminus DASD (Asp-Ala-Ser-Asp) sequence of the corresponding chains in TrK13-WT and appears to be involved in conferring stability to the overall hexameric assembly (shown as orange spheres in **Supplementary Figure 5**). To date, very few mutations are reported in the segments which seal the hexameric association. These possibly implicate that the assembly disrupting mutations could be unfavorable and are evolutionarily dissuaded. Further experiments are necessary to deliberately engineer mutations at the N-terminus of TrK13-WT to validate if mutations in these regions will derail PfKelch13 functioning and pathogen survival. If true, then this information could be a key to search for susceptible versus resistant parasites and the discovery of new range of molecules which may dock/bind to associating segment leading to assembly disruption.

The BTB domain of PfKelch13 shows highest similarity to KCTD17 [33]. Conventionally, BTB domains directly interact with the N-terminal domain of the cullin family protein Cul3 to constitute the Cullin-RING E3 ligase complexes (CRL3) together with the RING-domain protein Rbx1 and E2 ubiquitin-conjugating enzyme charged with ubiquitin [58]. Neddylation of Cul3 promotes a conformational change that culminates in efficient ubiquitination of the E3 bound substrate. While most of the BTB family proteins have a fused ligand-recognition domain to serve as substrate recognition subunit of the E3 ubiquitin ligase, in some cases proteins like Elongin C acts as an additional E3 ubiquitin ligase adaptor mediating the ubiquitination of substrates [59]. KCTD17 also exhibits E3 ligase activity and controls ciliogenesis by inducing degradation of trichoplein, although it lacks the corresponding downstream KRP regions [60]. Size-exclusion chromatography and SAXS data analyses have independently revealed that KCTD17 displays a closed pentameric oligomerization assembly to form a 5:5 heterodecameric geometry with Cul3[34]. Incidentally, KCTD17 lacks an additional ‘3-box’ motif found just C-terminus to the BTB domain in MATH-BTB and BTB-BACK-Kelch family E3 ligases and implicated in increasing the binding affinity to the N-terminus extensions in Cul3. The close proximity of the BTB domains in the KCTD17 pentamer are alleged to create additional Cul3-binding interface compensating for the lack of a 3-box, a model also supported by cryo-EM studies [61, 62]. Since PfKelch13 also lacks the 3-box motif, an association into a hexameric assembly, as evident from this study, may facilitate the creation of such additional sites for the cooperative binding of corresponding (six) copies of the Cul3 homolog in *P. falciparum*. Alternatively, the binding of PfCul3 may itself further stabilize the PfKelch13 hexamer and induce recruitment of the other homologs of CRL3 in *P. falciparum* to ensure efficient ubiquitination of substrates for their degradation. *P. falciparum* genome has two putative Cullin(like) proteins: PF3D7_0811000 and PF3D7_0629800, with PF3D7_0811000 exhibiting highest similarity with the *H. sapiens* Cul3 (data not shown). As of date, no association of PF3D7_0811000 has been detected in any immunoprecipitation or proximity biotinylation BioID studies [24, 25]. It remains to be established if PfKelch13 indeed functions as a substrate adaptor for CRL3 in *P. falciparum* and mediates ubiquitination of suitable substrates and if mutations in PfKelch13 influence Cullin binding or any subsequent downstream events. In only in a single study, it is reported that parasites with the dominant artemisinin-resistant C580Y mutation in PfKelch13 have higher levels of PfPI3K (a parasite protein, deemed as a substrate for PfKelch13) as compared to the wild-type. The C580Y mutation was speculated to have reduced or obliterated PfKelch13’s affinity towards PfPI3K to facilitate ubiquitin-mediated degradation. Resistant parasites thus displayed an increased abundance of PI3P that helped them overcome artemisinin-induced proteopathy [16, 26].

In the current study, we observed fine differences between the shape-assembly of TrK13-WT and the dominant artemisinin-resistant mutants TrK13-C580Y and TrK13-R539T, while the artemisinin-sensitive TrK13-A578S retained features similar to the TrK13-WT. However, in the absence of any resolved crystal structure for any PfKelch13 mutants, we are hesitant in over-interpreting our data due to limits in the resolution of the SAXS datasets and shape restoration or the uncertainty in the modeling routines. Nonetheless, the outcome of this study implies that specific mutations in PfKelch13 incorporate subtle conformational changes in its hexameric assembly. These might interfere in the formation of an active multi-subunit ubiquitination complex and/or influence the selective ubiquitination of specific substrates, either of which may be significant in ensuring survival of these parasites during artemisinin exposure. This work is a first major step in visualizing PfKelch13 and properties it holds. Overall, this study also paves the way for holistic intervention model which unravels new avenues for mapping the shape changes induced by other deleterious mutations in PfKelch13 and suggest ways to reverse these alterations in the native wild type PfKelch13 by small molecular interventions. Expression of other mutations in PfKelch13 and design of structurally-inspired resistant or rescue mutations forms the next challenge in our understanding of PfKelch13 which currently is the only confirmed marker for artemisinin-resistance. Until then, our results offer the community to address this aspect of malaria from their expertise alongside us.

## Materials and Methods

### Cloning, expression, and purification of recombinant PfKelch13 wild type and mutants from E. coli

The construct pPR-IBA101 (PfKelch13-WT) was generated by amplifying full-length *pfkelch13* (PF3D7_1343700) from *P. falciparum* 3D7 genomic DNA as template and using 101PfKelch13-F (5′-CAAATGGGAGACCTTATGGAAGGAGAAAAAGTAAAAACAAAAGCAAATAGTATCTCG-3′) and 101PfKelch13-R (5′-CCAAGCGCTGAGACCAGCAGCTATATTTGCTATTAAAACGGAGTGACCAAATCTGGG-3′) primers. The PCR product was digested with *BsaI* and cloned at corresponding sites in similarly digested pPR-IBA101 plasmid (IBA, Germany). The constructs pPR-IBA101 (PfKelch13-C580Y), pPR-IBA101 (PfKelch13-R539T) and pPR-IBA101 (PfKelch13-R539T) were generated by using overlapping PCR strategy. Briefly, the products of PCR1 using 101PfKelch13-F and C580Y-reverse (5′-TTATTATCAAAAGCAACATACATAGCTGATGATCTAGG-3′)/ R539T-reverse (5′-TGTACCGTTAGACGTAACACCACAATTATTTCTTCTAGGTATATTTAAATTAC-3′)/A578SR (5′-GCAACACACATAGATGATGATCTAGGGGTATTCAAAGGTGC-3′), and PCR2 using C580Y-forward (5′-CCTAGATCATCAGCTATGTATGTTGCTTTTGATAATAA-3′)/R539T-forward (5′-AGAAATAATTGTGGTGTTACGTCTAACGGTACAATTTATTGTATTGGGGGATATGATG-3′)/A578SF (5′-CCCTAGATCATCATCTATGTGTGTTGCTTTTGATAATAAAATTTATGTCATTGGT-3′) and 101PfKelch13-R were used as templates for the overlapping PCR3 to generate full-length *pfkelch13* (with residues conferring C580Y, R539T or A578S mutations in the protein sequence, respectively). The products of PCR3 were subsequently digested with *BsaI* and cloned into pPR-IBA101 to generate pPR-IBA101 (PfKelch13-C580Y), pPR-IBA101 (PfKelch13-R539T) and pPR-IBA101 (PfKelch13-A578S). Transformed *E. coli* DH10B colonies in each case were screened by restriction digestion and clones were confirmed by Sanger sequencing.

Confirmed clones were used to transform *E. coli* arabinose-inducible (AI) cells. For protein expression and purification, one liter of LB media with 100 µg/ml Ampicillin was used for each protein induction, which was then inoculated with overnight grown culture and kept at 37°C under shaking condition till they reached an O.D_600 nm_ of 0.6. Cultures were cooled to 22°C and induced with 1 mM isopropyl-1-thio-β-D-galactopyranoside (IPTG) and 0.2% Arabinose for overnight under shaking condition. Induced bacterial cells were harvested for 20 min at 6000 × *g* and the resulting pellet was resuspended in buffer W (100 mM Tris. HCl pH 8.0, 150 mM NaCl, 1 mM EDTA) containing 1 mM Phenylmethyl sulfonyl fluoride (PMSF) and 0.2 mg/ml lysozyme. Resuspended cells were incubated gently under rotating conditions at 4°C for 1 h and frozen at −80°C for overnight. At the next day, cells were then sonicated in ice for 4 min using a 30 s ‘on’ and 30 s ‘off’ pulse. The homogenate was centrifuged at 10,000 rpm at 4°C to remove insoluble debris, and the supernatant was collected into a 50 ml tube.

Recombinant TrK13-WT, TrK13-C580Y, TrK13-R539T and Trk13-WT were affinity purified from cellular lysate using StrepTactin XT Superflow resin (IBA, Germany). Briefly, soluble lysates were allowed to bind to resin overnight under rotating conditions at 4°C and then packed onto empty columns. The resin was washed extensively with at least 50 volumes of lysis buffer till no protein was detected in the flow through by Bradford’s method. Resin bound TrK13-WT, TrK13-C580Y, TrK13-R539T and TrK13-A578S were eluted with lysis buffer containing 50 mM biotin. Aliquots from eluted fractions were resolved by SDS-PAGE, pooled and dialyzed overnight at 4°C against PBS, pH 7.4 with at least three periodic changes. Protein samples were subsequently concentrated using Amicon 3-kDa concentrators (Millipore). The purity of each protein was detected by SDS-PAGE and western blotting. Purified proteins were stored at 4°C till further analyses.

### Generation of antibodies to PfKelch13

Custom antibodies were generated in rabbits against the peptide **C**EHRKRFDEERLRFL corresponding to amino acids 297-310 of PfKelch13 sequence (the cysteine at the N-terminus was added for conjugation to KLH) by Genscript Inc. The specificity of the antibody was confirmed by western blots against purified recombinant PfKelch13.

### Mass Spectrometry analyses and peptide quantitation

Mass spectrometry of purified TrK13-WT was performed at Wistar Proteomics Facility, Philadelphia, PA, USA. Protein bands of interest were excised from the gel, de-stained, alkylated with iodoacetamide, and digested with AspN or trypsin as previously described [16]. Digests were analyzed by LC-MS/MS using reverse phase capillary HPLC with a 75 µm nano-column interfaced on-line with a Thermo Electron LTQ OrbiTrap XL mass spectrometer. The mass spectrometer was operated in data dependent mode with full scans performed in the Orbitrap and parallel MS/MS analysis of the six most intense precursor ions in the linear ion trap. Resulting precursor masses and spectra were searched against a custom database using the Sequest search engine (Thermo Scientific) with the Proteomics Browser interface (provided by William Lane, Harvard Microchemistry and Proteomics Analysis Facility). The custom database consisted of a combination of the expected sequences with the various mutations of the proteins used in this experiment, the E. coli proteome (to provide a reasonable sized database), and expected common contaminants including keratins, trypsin, etc. A reversed sequence database was appended to the front of the forward database and used to estimate peptide false positive rates. Data were searched using partial tryptic specificity, a maximum of three missed cleavages, mass tolerance of 100 ppm, cysteine fixed as the carboxymethyl derivative, and dynamic methionine oxidation and N-terminal acetylation. Resulting data was filtered on 5ppm and dCn of 0.07. The false positive rate for peptide identifications using these database search and data filtering parameters was less than 3%.

### Western blotting

Western blots were performed using uninduced *E. coli* or IPTG-induced *E. coli* expressing TrK13-WT, TrK13-C580Y and TrK13-R539T using either commercial anti-STrEP (IBA, Germany) or custom-generated anti-PfKelch13 antibodies. Briefly, samples were solubilized in SDS sample buffer. Resolved by SDS-PAGE and transferred onto nitrocellulose membrane. The membrane was blocked with 5% skimmed milk in PBS, pH 7.4 for 1 h and incubated with the primary antibodies for overnight under shaking conditions at 4°C. After washing with PBS containing 0.05% Tween-20, the membrane was incubated with respective (anti-rabbit or anti-mouse) horseradish (HRP)-conjugated secondary antibodies. Blots were developed using chemiluminescence method.

### Small Angle X-ray Scattering (SAXS) experiments

Proteins were purified by size exclusion chromatography using Superdex200 10/300 GL column attached to AKTA purifier (GE Healthcare). Analytical gel filtration experiments were done using SHODEX Protein KW-800 GFC column (Showa-Denkko-KK, Japan) attached to a WATERS HPLC system. All SAXS experiments described here were done at SAXSpace Instrument with line collimation [63, 64]. CuKα X-rays generated from sealed tube source were used and scattering information was collected on 1D Mythen detector (Dectris). About 60 μL of WT and mutant proteins, and matched buffer were loaded in 1 mm thermostated quartz capillary. Initial experiments were done at 283 K, and later experimental temperature was varied from 283° to 353°C. At each temperature, SAXS data was acquired for 60 minutes of one frame. All programs employed for scattering data processing to primary analysis are reported in **Supplementary Table 3**. Shape parameters were analyzed using in-built modules in ATSAS 3.0.1 suite of programs. Intensity values at zero angles, I_0_ were estimated using Guinier approximation considering globular shape profile and estimation of Distance Distribution Function. These I_0_ values were compared with those from standard protein samples of lysozyme and gelsolin in EGTA buffer [63, 65]. Using restraints deduced from SAXS data analysis, chain-ensemble shape restoration protocol was utilized to construct solution shape of WT and mutants. Different symmetry options were considered, as reported earlier [66, 67] during shape restoration and the best convergence were seen for P6 symmetry. Some programs were run at online portal, and for others offline mode was used. Residue scale models were made using templates and energy optimization by Swiss Model Server. Using single chain of model of TrK13-WT and mutants and placement by using P6 symmetry using SAXS data as reference by SASREF program. All images for molecular shapes were prepared using UCSF Chimera (*https://www.cgl.ucsf.edu/chimera/*) and PyMol programs. Some graphs shown in this work were plotted using OriginLab version5 software (*www.originlab.com*).

## Acknowledgements

The authors would like to thank Drs. David W Speicher and Kaye D Speicher at the Wistar Proteomics Core Facility, The Wistar Institute, Philadelphia, PA, USA for their help in LC-MS/MS and analyses. The authors would also like to thank Prof. Kasturi Haldar and Dr. Niraja Suresh, University of Notre Dame, USA for their helpful suggestions during the designing of the study and assembly of the manuscript. This work was supported by funding from the Science and Engineering Research Board, Department of Science and Technology, Government of India (DST ECR/2015/000387), Department of Biotechnology Ramalingaswami Re-entry Fellowship (BT/HRD/35/02/ 2006) and University for Potential Excellence–II (Project ID 245) (S.B.). N.G. is a recipient of research fellowship from the Council of Scientific and Industrial Research, Government of India.

## Author contributions

N.G., K.D., N.K., A.M., A. and S.B. designed, performed, and interpreted the experimental work; A.M. helped in the experiments; A. and S.B. provided key intellectual insight into aspects of this study and wrote the manuscript; and all the authors commented on the manuscript.

## Additional Information

The coordinates of models generated in this study are available on request for academic purposes.

## List of Supplementary files

**Supplementary Figure Legends**

**Supplementary Tables 1-3**

**Supplementary Figures 1-5**

