## Supplemental data for "*Plasmodium falciparum* Kelch13 and its artemisinin-resistant mutants assemble as hexamers in solution: a SAXS data driven shape restoration study"

**Affiliations:**
[a]Special Centre for Molecular Medicine, Jawaharlal Nehru University, New Delhi-110067, India
[b]Council of Scientific and Industrial Research-Institute of Microbial Technology, Chandigarh-160036, India

*Corresponding author

**Address for correspondence:** Souvik Bhattacharjee; 3D-FF, Special Centre for Molecular Medicine, Jawaharlal Nehru University, New Delhi-110067, INDIA; Phone: +919810125616, +911126738974,

***List of Supplementary files***

**Supplementary Figure Legends**

**Supplementary Tables 1-3**

**Supplementary Figures 1-5**

**Supplementary Figure 1. LC-MS/MS analyses of TrK13-WT after trypsin or AspN digestion. a.** Table showing the sequences of detected peptides, their counts and position within the amino acid sequence of PfKelch13-strep after Trypsin digestion. Deamidation and oxidation of specific residues are highlighted in red and green, respectively. **b.** LC-MS/MS analyses of TrK13-WT after AspN digestion indicating the positions and lengths of detected peptides. The most N-terminus peptide detected is KFSTVNNVN. **c.** Table showing the sequences of detected peptides, their counts and position within the amino acid sequence of PfKelch13-strep after AspN digestion. Oxidation of specific amino acid residues are highlighted in green.

**Supplementary Figure 2. SAXS data of TrK13-WT. a.** Double log plot showing SAXS datasets from three different concentrations of TrK13-WT. SAXS profiles from freshly purified TrK13-WT at the concentrations of 1.4, 0.8 and 0.25 mg/ml are plotted as red, magenta and blue symbols, respectively. The lower inset shows the dimensionless Kratky plot of the dataset at 1.4 mg/ml. Right upper and lower plots show the Guinier analysis and residuals for the fit (red line) for the same dataset.

**Supplementary Figure 3. VTSAXS profiles of TrK13-WT and mutants.** VTSAXS profiles from samples of purified TrK13-WT (**a**), TrK13-R539T (**b**), TrK13-C580Y (**c**) and TrK13-A578S (**d**). Left panels show the datasets at different temperatures for samples at TrK13-WT (1.4 mg/ml), TrK13-R539T (1.1 mg/ml), TrK13-C580Y (1.3 mg/ml) and TrK13-A578S (0.9 mg/ml)]. Right panel shows the Kratky plot of the datasets plotted on left. The temperatures of the dataset are as indicated. All exposures were for 60 minutes each.

**Supplementary Figure 4. TrK13-WT structure prediction, domain assignment, fragmentation and ligation.** Domains in TrK13-WT abstracted for sequence similarity-based template search combined with energy optimization and later used for positioning in space to generate models which would agree with SAXS data-based shape for the hexameric assemblies. **a.** Predicted structure of TrK13-WT based on the amino acid sequence using Phyre2 server (*www.sbg.bio.ic.ac.uk/phyre2*) and rainbow colored in PyMol. The strep-tag at the C-terminus is colored grey. **b.** Sequence of TrK13-WT highlighting sequence of each domains by independent colors and the corresponding amino acid residues. **c.** Representative chain of TrK13-WT model showing different orientations of segments stitched together after their relative positioning in space using SWISS Modeler server (*https://swissmodel.expasy.org/*).

**Supplementary Figure 5. Location of N-terminus mutation in PfKelch13 with respect to TrK13-WT.** Positions of K189T (red), I205T (magenta), E252Q (yellow) and R255K (green) mutations in PfKelch13 are shown as spheres in the representative TrK13-WT hexamer and two different orientations. The amino acid residues DASD motif, where the individual chains of the hexamer coalesce together, are indicated by orange spheres.

**Supplementary Table 1. Potential cleavage sites in PfKelch13-STrEP cleaved by proteases or chemicals.** (*https://web.expasy.org/peptide_cutter/*).

**Supplementary Table 2. Epidemiological information on various mutations in PfKelch13.** Table showing the various PfKelch13 mutations, residue position in TrK13-WT, country and region of origin, location of the mutation and assignment of resistance values based on the range of $xPC_{1/2}$ [30]

**Supplementary Table 3. SAXS data collection and scattering parameters**

**Supplementary Table 1: Potential cleavage sites in PfKelch13-STrEP cleaved by proteases or chemicals (*https://web.expasy.org/peptide_cutter/*)**

**Sequence of full length PfKelch13-STrEP: Blue text, PfKelch13; Red, STrEP tag**

MEGEKVKTKANSISNFSMTYDRESGGNSNSDDKSGSSSENDSNSFMNLTSDKNEKTENNSFLLNNSSYGNVKDSLLE
SIDMSVLDSNFDSKKDFLPSNLSRTFNNMSKDNIGNKYLNKLLNKKKDTITNENNNINHNNNNNNLTANNITNNLIN
NNMNSPSIMNTNKKENFLDAANLINDDSGLNNLKKFSTVNNVNDTYEKKIIETELSDASDFENMVGDLRITFINWLK
KTQMNFIREKDKLFKDKKELEMERVRLYKELENRKNIEEQKLHDERKKLDIDISNGYKQIKKEKEEHRKRFDEERLR
FLQEIDKIKLVLYLEKEKYYQEYKNFENDKKKIVDANIATETMIDINVGGAIFETSRHTLTQQKDSFIEKLLSGRHH
VTRDKQGRIFLDRDSELFRIILNFLRNPLTIPIPKDLSESEALLKEAEFYGIKFLPFPLVFCIGGFDGVEYLNSMEL
LDISQQCWRMCTPMSTKKAYFGSAVLNNFLYVFGGNNYDYKALFETEVYDRLRDVWYVSSNLNIPRRNNCGVTSNGR
IYCIGGYDGSSIIPNVEAYDHRMKAWVEVAPLNTPRSSAMCVAFDNKIYVIGGTNGERLNSIEVYEEKMNKWEQFPY
ALLEARSSGAAFNYLNQIYVVGGIDNEHNILDSVEQYQPFNKRWQFLNGVPEKKMNFGAATLSDSYIITGGENGEVL
NSCHFFSPDTNEWQLGPSLLVPRFGHSVLIANIAAGLSAWSHPQFEKGGGSGGGSGGGSWSHPQFEK

| Name of the Enzyme | Number of Cleavages | Position of Cleavage Sites |
|---|---|---|
| Arg-C-Proteinase | 31 | 22 101 223 239 255 257 265 277 299 301 306 308 365 383 388 393 398 404 411 471 513 515 528 529 539 561 575 597 622 659 716 |
| Asp-N endopeptidase | 46 | 20 30 31 40 50 72 79 84 88 92 108 124 172 179 180 197 210 213 220 241 246 274 280 282 302 313 336 342 352 372 388 396 398 420 451 463 500 511 515 546 558 583 640 647 679 701 |
| Asp-N endopeptidase + N-terminal Glu | 106 | 1 3 20 22 30 31 38 40 50 53 56 72 76 79 84 88 92 108 124 129 168 172 179 180 197 200 205 207 210 213 215 220 239 241 246 249 251 253 260 262 268 269 274 275 280 282 293 295 296 302 303 304 311 313 322 324 329 334 336 342 348 352 361 372 376 388 396 398 400 420 423 425 430 432 451 454 460 463 500 506 508 511 515 546 555 558 566 583 595 601 604 605 611 619 640 642 647 650 667 679 687 690 701 704 738 758 |
| BNPS-Skatole | 9 | 229 470 518 565 611 660 706 733 753 |
| CNBr | 18 | 1 18 46 81 106 157 163 218 235 253 351 460 472 476 562 579 608 671 |
| Chymotrypsin-high specificity (C-term to [FYW], not before P) | 75 | 16 20 45 61 68 88 94 103 115 171 190 200 215 226 229 237 245 259 288 302 309 321 327 328 331 334 361 375 395 403 409 434 435 439 446 451 456 470 482 483 491 493 495 500 502 506 511 518 519 541 546 558 565 583 588 604 611 616 628 630 635 653 656 660 662 673 682 698 699 706 717 733 738 753 758 |
| Chymotrypsin-low specificity (C-term to [FYWML], not before P) | 166 | 1 16 18 20 45 46 48 61 62 63 68 75 76 81 84 88 94 99 103 106 115 116 119 120 136 143 152 157 163 171 172 177 184 187 190 200 209 215 218 222 226 229 230 235 237 244 245 251 253 258 259 262 273 280 288 298 302 307 309 310 318 320 321 322 327 328 331 334 351 361 366 368 375 379 380 384 385 395 396 402 403 407 409 410 414 422 428 429 434 435 439 444 446 451 456 457 460 462 463 470 472 476 482 483 488 491 492 493 495 500 502 505 506 511 514 518 519 524 541 546 558 560 562 565 571 579 583 588 598 604 608 611 616 618 619 628 630 631 635 644 647 653 656 660 662 663 671 673 678 682 693 697 698 699 706 708 712 713 717 719 722 730 733 738 753 758 |

| | | |
|---|---|---|
| Clostripain | 31 | 22 101 223 239 255 257 265 277 299 301 306 308 365 383 388 393 398 404 411 471 513 515 528 529 539 561 575 597 622 659 716 |
| Formic acid | 46 | 21 31 32 41 51 73 80 85 89 93 109 125 173 180 181 198 211 214 221 242 247 275 281 283 303 314 337 343 353 373 389 397 399 421 452 464 501 512 516 547 559 584 641 648 680 702 |
| Glutamyl endopeptidase | 60 | 2 4 23 39 54 57 77 130 169 201 206 208 216 240 250 252 254 261 263 269 270 276 294 296 297 304 305 312 323 325 330 335 349 362 377 401 424 426 431 433 455 461 507 509 556 567 596 602 605 606 612 620 643 651 668 688 691 705 739 759 |
| Hydroxylamine | 5 | 286 537 594 664 689 |
| Iodosobenzoic acid | 9 | 229 470 518 565 611 660 706 733 753 |
| LysC | 64 | 5 7 9 33 52 55 72 91 92 108 114 118 122 123 124 167 168 188 189 202 203 231 232 241 243 246 248 249 260 266 272 278 279 289 292 293 295 300 315 317 324 326 332 338 339 340 372 378 390 420 430 438 479 480 503 563 586 607 610 658 669 670 740 760 |
| LysN | 64 | 4 6 8 32 51 54 71 90 91 107 113 117 121 122 123 166 167 187 188 201 202 230 231 240 242 245 247 248 259 265 271 277 278 288 291 292 294 299 314 316 323 325 331 337 338 339 371 377 389 419 429 437 478 479 502 562 585 606 609 657 668 669 739 759 |
| NTCB (2-nitro-5-thiocyanobenzoic acid) | 7 | 446 468 472 531 541 579 695 |
| Pepsin (pH1.3) | 134 | 15 16 44 45 47 48 60 61 62 63 75 76 83 84 87 88 95 98 99 102 115 118 119 142 143 151 152 171 172 176 177 183 184 186 187 189 208 209 214 215 221 222 226 229 230 236 237 244 258 261 272 273 307 309 318 320 321 322 333 360 361 375 378 379 394 396 401 402 403 407 408 409 410 421 427 428 429 433 434 438 443 445 446 450 451 456 457 461 462 463 483 487 488 490 491 492 494 495 504 506 514 523 524 570 582 583 598 614 617 618 619 627 628 630 631 647 655 662 663 673 677 678 692 693 697 698 707 712 717 722 729 730 738 758 |
| Pepsin (pH>2) | 184 | 15 16 19 20 44 45 47 48 60 61 62 63 67 68 75 76 83 84 87 88 95 98 99 102 114 115 118 119 142 143 151 152 171 172 176 177 183 184 186 187 189 199 200 208 209 214 215 221 222 226 228 229 230 236 237 244 258 261 272 273 287 288 307 309 318 320 321 322 327 330 331 333 360 361 375 378 379 394 396 401 402 403 407 408 409 410 421 427 428 429 433 434 435 438 443 445 446 450 451 455 456 457 461 462 463 469 470 483 487 488 490 491 492 493 494 495 499 500 501 502 504 506 510 511 514 518 519 523 524 540 545 546 557 558 564 570 582 583 587 598 603 604 610 611 614 615 617 618 619 627 628 629 630 631 634 635 647 652 655 662 663 673 677 678 681 682 692 693 697 698 705 706 707 712 717 722 729 730 732 733 738 752 753 758 |
| Proline-endopeptidase [*] | 2 | 736 756 |
| Proteinase K | 343 | 2 4 6 8 10 13 16 19 20 23 39 45 48 49 54 56 57 61 62 63 68 71 75 76 77 79 83 84 88 94 95 99 102 103 111 115 116 119 120 126 127 128 130 134 143 144 145 148 149 152 153 162 165 169 171 172 174 175 177 178 184 |

| | | |
|---|---|---|
| | | 187 190 192 193 196 199 200 201 204 205 206 207 208 209 212 215 216 219 222 224 225 226 227 229 230 233 237 238 240 244 245 250 251 252 254 256 258 259 261 262 263 268 269 270 273 276 280 282 284 288 291 294 296 297 302 304 305 307 309 310 312 313 316 318 319 320 321 322 323 325 327 328 330 331 334 335 341 342 344 346 347 348 349 350 352 354 356 359 360 361 362 363 367 368 369 375 376 377 379 380 386 387 394 395 396 401 402 403 405 406 407 409 410 414 415 416 418 422 424 426 427 428 429 431 432 433 434 435 437 439 440 442 444 445 446 448 451 454 455 456 457 461 462 463 465 470 474 478 481 482 483 486 487 488 491 492 493 494 495 500 502 504 505 506 507 508 509 510 511 514 517 518 519 520 524 526 534 535 540 541 543 546 551 552 555 556 557 558 564 565 566 567 568 569 571 573 578 581 582 583 587 588 589 590 593 596 598 601 602 603 604 605 606 611 612 614 616 617 618 619 620 621 626 627 628 630 631 634 635 636 637 640 643 646 647 650 651 653 656 660 662 663 666 668 673 675 676 677 678 682 683 684 685 688 691 692 693 698 699 703 705 706 708 712 713 714 717 721 722 723 724 726 727 728 730 732 733 738 739 753 758 759 |
| Staphylococcal peptidase I | 56 | 2 4 23 39 54 57 77 130 169 201 206 208 216 240 250 252 254 261 263 269 276 294 296 304 312 323 325 330 335 349 362 377 401 424 426 431 433 455 461 507 509 556 567 596 602 605 612 620 643 651 668 688 691 705 739 759 |
| Thermolysin | 192 | 5 9 12 15 17 44 45 47 60 61 62 70 74 75 78 82 83 87 98 102 105 110 115 118 119 126 133 142 144 147 151 152 156 161 162 170 171 174 176 177 183 186 189 192 195 203 204 217 218 223 225 226 229 234 236 237 243 244 255 257 267 272 279 290 301 306 308 309 315 317 318 319 321 333 340 341 345 346 350 351 355 358 359 360 367 374 375 378 379 385 393 394 395 402 404 405 406 408 409 413 427 428 436 438 443 444 445 447 450 453 456 459 462 471 475 480 482 485 486 487 490 491 493 494 503 504 505 513 519 523 533 539 542 550 554 561 563 565 570 577 578 580 581 582 586 588 589 597 600 607 616 617 618 625 626 627 630 633 635 636 639 645 646 649 655 661 662 670 672 674 675 677 682 683 692 697 698 707 711 712 716 720 721 722 723 725 726 727 729 731 737 757 |
| Thrombin | 1 | 716 |
| Trypsin | 95 | 5 7 9 22 33 52 55 72 91 92 101 108 114 118 122 123 124 167 168 188 189 202 203 223 231 232 239 241 243 246 248 249 255 257 260 265 266 272 277 278 279 289 292 293 295 299 300 301 306 308 315 317 324 326 332 338 339 340 365 372 378 383 388 390 393 398 404 411 420 430 438 471 479 480 503 513 515 528 529 539 561 563 575 586 597 607 610 622 658 659 669 670 716 740 760 |
| These chosen enzymes do not cut: | | |

Caspase1

Caspase10

Caspase2

Caspase3

Caspase4

Caspase5

Caspase6

Caspase7

Caspase8

Caspase9

Enterokinase

Factor Xa

GranzymeB

Tobacco etch virus protease

**Supplementary Table 2: PfKelch13 mutants selected in this study.** Table showing the various PfKelch13 mutations, residue position in TrK13-WT, country and region of origin, location of the mutation and assignment of resistance values based on the range of $xPC_{1/2}$ [30].

| S. No. | Kelch mutation | Numbering with respect to TrK13-WT | Country | Location | Mutation site | $xPC_{1/2}$ values [30]. <1.39 = Blank/sensitive, 1.4-1.69 = +/Low resistance, 1.7-1.99 = ++/Medium resistance, ≥2 = +++/High Resistance |
|---|---|---|---|---|---|---|
| 1 | N458Y | N270Y | Thailand, | Thai Western border, | Propeller 1 | (2.5) +++ |
| 2 | P527H | P339H | Thailand, | Thai Western border, | Propeller 2 | (1.7) ++ |
| 3 | Y493H | Y305H | Viet Nam, Cambodia, Viet Nam | Tra Leng, Pursat, Bin Phuoc | Propeller 2 | (2.7) +++ |
| 4 | K189N | K1N | Bangladesh, Cambodia | Ramu, Preah Vihear | N-terminus | (1.1) sensitive/0 |
| 5 | R539T | R351T | Cambodia, Cambodia, Viet Nam | Pailin, Pursat, Bin Phuoc | Propeller 2 | (2.1) +++ |
| 6 | C580Y | C392Y | Cambodia, Cambodia, Cambodia, Thailand, Thailand, Thailand, Viet Nam, Cambodia | Pailin, Pursat, Ratanakiri, Ranong, Thai Western border, Srisaket, Bin Phuoc, Preah Vihear | Region between propeller 3 and 4 | (2.3) +++ |
| 7 | P553L | P365L | Cambodia, Viet Nam | Preah Vihear, Bin Phuoc | Propeller 3 | (2.2) +++ |
| 8 | D584V | D396V | Cambodia | Pursat | Region between propeller 3 and 4 | Not available |
| 9 | K189T | K1T | DRC | Kinshasa | N-terminus | (1.1) sensitive/0 |
| 10 | I543T | I355T | Lao PDR, Viet Nam | Attapeu, Bin Phuoc | Propeller 3 | (average of 2.1 and 2.8 = 2.45) +++ |
| 11 | E252Q | E64Q | Myanmar, Thailand | Shwe Kyin, Thai Western border | Helical region | (1.5) + |
| 12 | P441L | P253L | Myanmar, Thailand | Shwe Kyin, Thai Western border | Uncharacterized region | (2.2) +++ |
| 13 | F446I | F307I | Myanmar, Thailand | Shwe Kyin, Thai Western border | Propeller 1 | (1.5) + |
| 14 | P574L | P386L | Thailand | Ranong | Propeller 3 | (average of 1.8 and 2 = 1.9) ++ |
| 15 | G538V | G350V | Thailand | Thai Western border | Propeller 2 | (1.9) ++ |
| 16 | R561H | R373H | Thailand | Thai Western border | Propeller 3 | (2.2) +++ |
| 17 | S621F (S623F)? | S435F | Thailand | Tak provinence | Region between propeller 4 and 5 | Not available |
| 18 | A675V | A487V | Thailand | Mae Hong Son | Propeller 5 | (2.2) +++ |
| 19 | A481V | A293V | Thailand | Mae Hong Son | Propeller 1 | (1.6) + |
| 20 | M476I | M288I | Thailand | Mae Hong Son | Propeller 1 | (2.0) +++ |
| 21 | N525D | N337D | Thailand | Mae Hong Son | Propeller 2 | Not available |
| 22 | N572I | N384I | Thailand | Mae Hong Son | Propeller 3 | Not available |
| 23 | F495L | F307L | Thailand | Mae Hong Son | Propeller 3 | Not available |
| 24 | E567D | E379D | Thailand | Kanchanaburi | Propeller 3 | Not available |
| 25 | R575K | R387K | Thailand | Kanchanaburi | Region between propeller 3 and 4 | Not available |
| 26 | G449A | G261A | Thailand | Ubon | Propeller 1 | (1.9) ++ |
| 27 | V566I | V378I | Africa/Ghana | | Propeller 3 | Not available |
| 28 | G533A | G345A | India, Myanmar | | Propeller 2 | Not available |
| 29 | A578S | A390S | Africa | | Between propeller 3 and 4 | Not available |
| 30 | V568G | V380G | Viet Nam | Thuy Nhien | Propeller 3 | (2.7) +++ |

**Supplementary Table 3. SAXS data collection and scattering parameters**

| | |
|---|---|
| **Data-collection parameters** | |
| Instrument | SAXSpace (Anton Paar) |
| Beam geometry | 10 mm slit |
| Wavelength (Å) | 1.5418 |
| Desmearing Software | Done using SAXSquant |
| Exposure time for each sample/buffer | 60 minutes |
| q range ($nm^{-1}$) | 0.08–6.00 |
| Temperature (K) | 298-353 |
| **Data analysis and modeling programs employed** | |
| Beam Position Correction | SAXStreat |
| Primary data reduction | SAXSquant |
| Data processing | PRIMUS QT (ATSAS 3.0.1) |
| Ab initio Shape Restoration | GASBOR P6 570 DR Ten Models Each [Online] |
| Validation and averaging | DAMAVER [Offline] |
| Computation of model intensities | CRYSOL [Offline] |
| Spatial positioning of domains in space using SAXS | SASREF [Offline] |
| Plotting of data or fits | ATSAS Interface |
| Plotting of P(r) curves | Origin 5.0 |
| Three-dimensional graphics representations | PyMOL; UCSF Chimera 1.12 |
