## Supplemental Figures for "*Plasmodium falciparum* Kelch13 and its artemisinin-resistant mutants assemble as hexamers in solution: a SAXS data driven shape restoration study"

### Supplementary Figure 1

## a

**Legends:**
■ +0.98 : Deamidation ■ +15.99 : Oxidation

| Peptide: | Count: | Position: | Peptide: | Count: | Position: |
|---|---|---|---|---|---|
| ALFETEVYDR | 1 | 504-513 | LDIDISNGYK | 1 | 280-289 |
| AWVEVAPLNTPR | 2 | 564-575 | LGPSLLVPR | 1 | 708-716 |
| CIGGFDGVEYLNSMELLDISQQCWR | 2 | 447-471 | LNSIEVYEEK | 2 | 598-607 |
| CIGGFDGVEYLNSMELLDISQQCWR | 1 | 447-471 | LNSIEVYEEK | 3 | 598-607 |
| CVAFDNK | 1 | 580-586 | LRDVWYVSSNLNIPR | 1 | 514-528 |
| DLSESEALLK | 5 | 421-430 | LTIPIPK | 1 | 414-420 |
| DVWYVSSNLNIPR | 4 | 516-528 | MCTPMSTK | 3 | 472-479 |
| EAEFYGIK | 4 | 431-438 | MCTPMSTK | 1 | 472-479 |
| ELEMER | 2 | 250-255 | MCTPMSTK | 2 | 472-479 |
| FGHSVL | 1 | 717-722 | MCTPMSTK | 2 | 472-479 |
| FGHSVLIAN | 1 | 717-725 | MNKWEQFPYALLEAR | 1 | 608-622 |
| FGHSVLIANIAAGL | 1 | 717-730 | NFLYVFGGNNYDYK | 1 | 490-503 |
| FGHSVLIANIAAGLSAWSHPQFEK | 3 | 717-740 | NNCGVTSNGR | 1 | 530-539 |
| FGHSVLIANIAAGLSAWSHPQFEK | 16 | 717-740 | NNCGVTSNGR | 2 | 530-539 |
| FLPFPLVF | 1 | 439-446 | NPLTIPIPK | 3 | 412-420 |
| FLPFPLVFC | 1 | 439-447 | NVNDTYEK | 2 | 195-202 |
| FLQEIDKIK | 1 | 309-317 | PLTIPIPK | 1 | 413-420 |
| FSTVNNVNDTYEK | 1 | 190-202 | QIYVVGGIDNEHNILDSVEQYQPFNK | 1 | 633-658 |
| GGGSGGGSGGGSWSHPQFEK | 3 | 741-760 | RWQFLNGVPEK | 2 | 659-669 |
| GSGGGSGGGSWSHPQFEK | 1 | 743-760 | RWQFLNGVPEK | 1 | 659-669 |
| IAAGLSAWSHPQFEK | 5 | 726-740 | SAWSHPQFEK | 1 | 731-740 |
| IANIAAGLSAWSHPQFEK | 1 | 723-740 | SCHFFSPDTNEWQLGPSLLVPR | 1 | 695-716 |
| IFLDRDSELFR | 1 | 394-404 | SMELLDISQQCWR | 1 | 459-471 |
| IIETELSDASDFENMVGDLR | 2 | 204-223 | SSAMCVAFDNK | 1 | 576-586 |
| IIETELSDASDFENMVGDLR | 2 | 204-223 | SSGAAFNYLNQIYVVGGIDNEHNILDSVEQYQPFNK | 1 | 623-658 |
| IIETELSDASDFENMVGDLR | 1 | 204-223 | SSGAAFNYLNQIYVVGGIDNEHNILDSVEQYQPFNK | 1 | 623-658 |
| IIETELSDASDFENMVGDLR | 6 | 204-223 | SSGAAFNYLNQIYVVGGIDNEHNILDSVEQYQPFNK | 1 | 623-658 |
| IILNFLR | 1 | 405-411 | SSGAAFNYLNQIYVVGGIDNEHNILDSVEQYQPFNK | 1 | 623-658 |
| IILNFLR | 1 | 405-411 | STVNNVNDTYEK | 1 | 191-202 |
| ISQQCWR | 1 | 465-471 | SWSHPQFEK | 1 | 752-760 |
| ITFINWLK | 1 | 224-231 | TQMNFIR | 1 | 233-239 |
| IYCIGGYDGSSIIPNVEAYDHR | 2 | 540-561 | TQMNFIR | 1 | 233-239 |
| IYCIGGYDGSSIIPNVEAYDHR | 3 | 540-561 | TQMNFIR | 1 | 233-239 |
| IYVIGGTNGER | 4 | 587-597 | VFGGNNYDYK | 1 | 494-503 |
| IYVIGGTNGER | 1 | 587-597 | VNNVNDTYEK | 1 | 193-202 |
| KFSTVNNVNDTYEK | 1 | 189-202 | VSSNLNIPR | 1 | 520-528 |
| KFSTVNNVNDTYEK | 1 | 189-202 | VVGGIDNEHNILDSVEQYQPFNK | 1 | 636-658 |
| KFSTVNNVNDTYEKK | 1 | 189-203 | VVGGIDNEHNILDSVEQYQPFNK | 1 | 636-658 |
| KKLDIDISNGYK | 2 | 278-289 | WEQFPY | 1 | 611-616 |
| KLDIDISNGYK | 4 | 279-289 | WEQFPYALLEAR | 6 | 611-622 |
| KLDIDISNGYK | 2 | 279-289 | WQFLNGVPEKK | 3 | 660-670 |
| KTQMNFIR | 1 | 232-239 | WQFLNGVPEKK | 1 | 660-670 |
| LDIDISNGYK | 4 | 280-289 | | | |

### Supplementary Figure 1 (contd.)

**b**

MEGEKVKTKA NSISNFSMTY DRESGGNSNS DDKSGSSSEN DSNSFMNLTS DKNEKTENNS FLLNNSSYGN VKDSLLESID MSVLDSNFDS
KKDFLPSNLS RTFNNMSKDN IGNKYLNKLL NKKKDTITNE NNNINHNNNN NNLTANNITN NLINNNMNSP SIMNTNKKEN FLDAANLIND
DSGLNNLKKF STVNNVNDTY EKKIIETELS DASDFENMVG DLRITFINWL KKTQMNFIRE KDKLFKDKKE LEMERVRLYK ELENRKNIEE
QKLHDERKKL DIDISNGYKQ IKKEKEEHRK RFDEERLRFL QEIDKIKLVL YLEKEKYYQE YKNFENDKKK IVDANIATET MIDINVGGAI
FETSRHTLTQ QKDSFIEKLL SGRHHVTRDK QGRIFLDRDS ELFRIILNFL RNPLTIPIPK DLSESEALLK EAEFYGIKFL PFPLVFCIGG
FDGVEYLNSM ELLDISQQCW RMCTPMSTKK AYFGSAVLNN FLYVFGGNNY DYKALFETEV YDRLRDVWYV SSNLNIPRRN NCGVTSNGRI
YCIGGYDGSS IIPNVEAYDH RMKAWVEVAP LNTPRSSAMC VAFDNKIYVI GGTNGERLNS IEVYEEKMNK WEQFPYALLE ARSSGAAFNY
LNQIYVVGGI DNEHNILDSV EQYQPFNKRW QFLNGVPEKK MNFGAATLSD SYIITGGENG EVLNSCHFFS PDTNEWQLGP SLLVPRFGHS
VLIANIAAGL SAWSHPQFEK GGGSGGGSGG GSWSHPQFEK

**c**

**Legends:**
+15.99 : Oxidation

| Peptide: | Count: | Position: | Peptide: | Count: | Position: |
|---|---|---|---|---|---|
| DEERLRFLQEI | 2 | 303-313 | DSFIEKLLSGRHHVTR | 8 | 373-388 |
| DGSSIIPNVEAY | 2 | 547-558 | DSVEQYQPFNKRWQFL | 2 | 648-663 |
| DHRMKAWVEVAPLNTPRSSAM | 1 | 559-579 | DTNEWQLGPSLLVPRFGHSVLIA | 1 | 702-724 |
| DHRMKAWVEVAPLNTPRSSAMCVAF | 1 | 559-583 | DVWYVSSNLNIPRR | 2 | 516-529 |
| DHRMKAWVEVAPLNTPRSSAMCVAF | 1 | 559-583 | DVWYVSSNLNIPRRNNCGVTS | 1 | 516-536 |
| DHRMKAWVEVAPLNTPRSSAMCVAF | 2 | 559-583 | DYKALFETEVY | 3 | 501-511 |
| DHRMKAWVEVAPLNTPRSSAMCVAF | 2 | 559-583 | EARSSGAAFNYL | 1 | 620-631 |
| DIDISNGYKQIKKEKEEHRKRF | 1 | 281-302 | EARSSGAAFNYLNQIY | 1 | 620-635 |
| DINVGGAIF | 1 | 353-361 | EARSSGAAFNYLNQIYVVGGIDNEHNIL | 2 | 620-647 |
| DINVGGAIFETSRHTLTQQK | 3 | 353-372 | EKGGGSGGGSGGGSWSHPQFEK | 1 | 739-760 |
| DISQQCWRMCTPMSTKKAYFGSAVLNNFLYVFGGNNY | 1 | 464-500 | EKKIIETELS | 1 | 201-210 |
| DKIKLVLY | 1 | 314-321 | ELEMERVRLYKELENRKNIE | 1 | 250-269 |
| DKIKLVLYL | 1 | 314-322 | EQKLHDERKKL | 1 | 270-280 |
| DKIKLVLYLEKEKYYQEYKNFEN | 2 | 314-336 | EVLNSCHFFSP | 1 | 691-701 |
| DKKELEMERVRLYKELENRKNIE | 5 | 247-269 | KFSTVNNVN | 1 | 189-197 |
| DKQGRIFL | 1 | 389-396 | LEKEKYYQEYKNFEN | 1 | 322-336 |
| DLRITFINWLKKTQMNFIREK | 2 | 221-241 | MNFIREK | 1 | 235-241 |
| DLRITFINWLKKTQMNFIREK | 3 | 221-241 | NIEEQKLH | 1 | 267-274 |
| DSFIEKLL | 1 | 373-380 | QIYVVGGIDNEHNIL | 1 | 633-647 |

**Supplementary Figure 2**

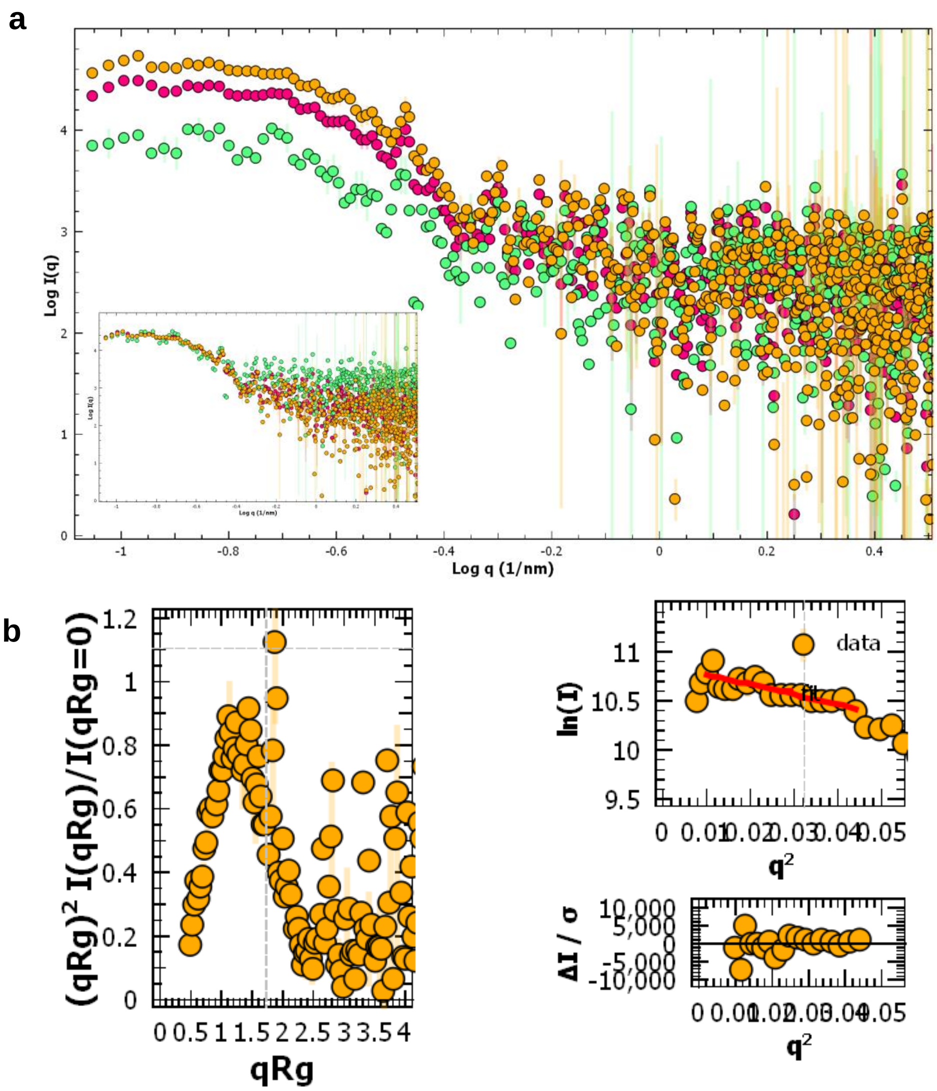

### Supplementary Figure 3

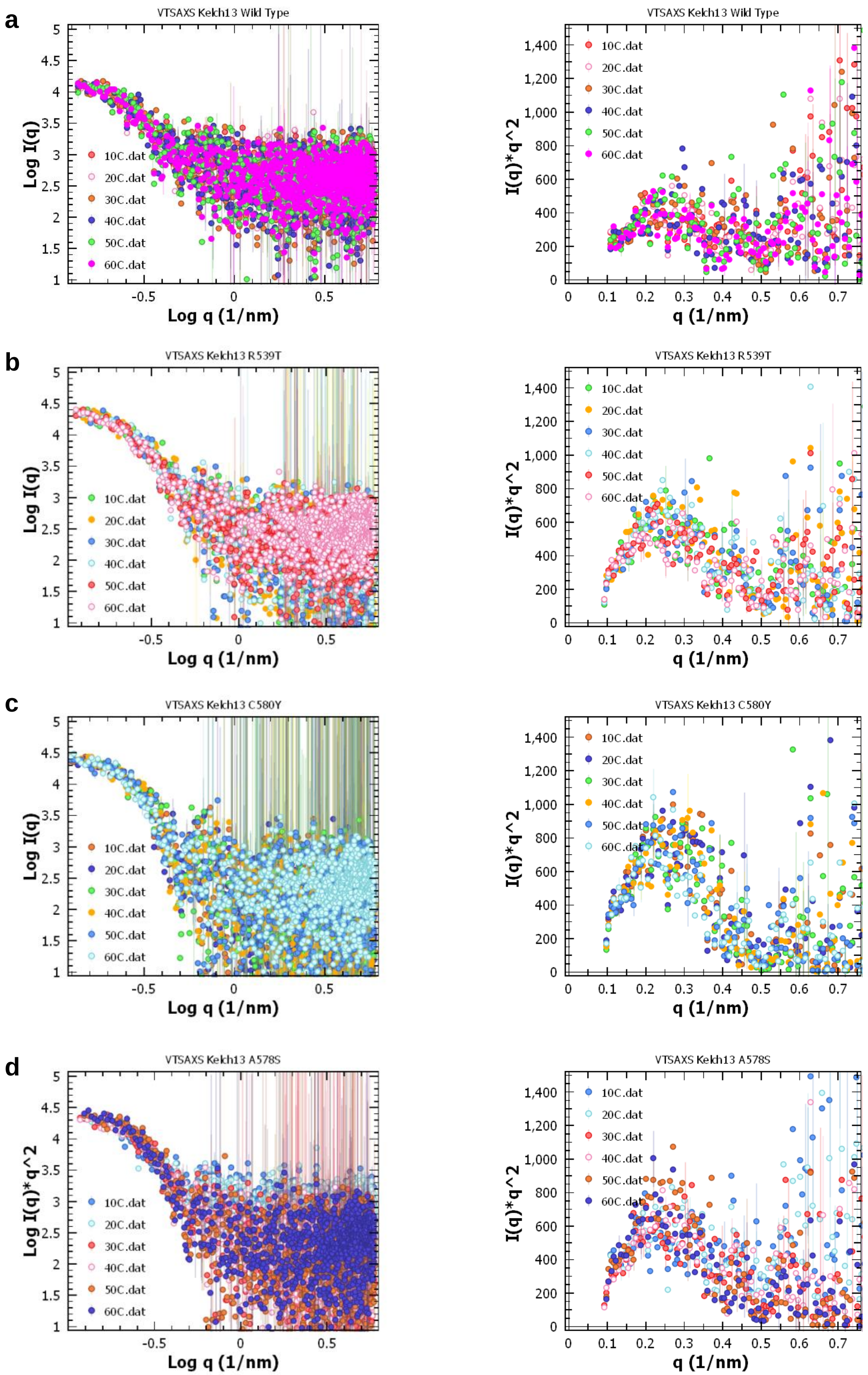

#### Supplementary Figure 4

**a**

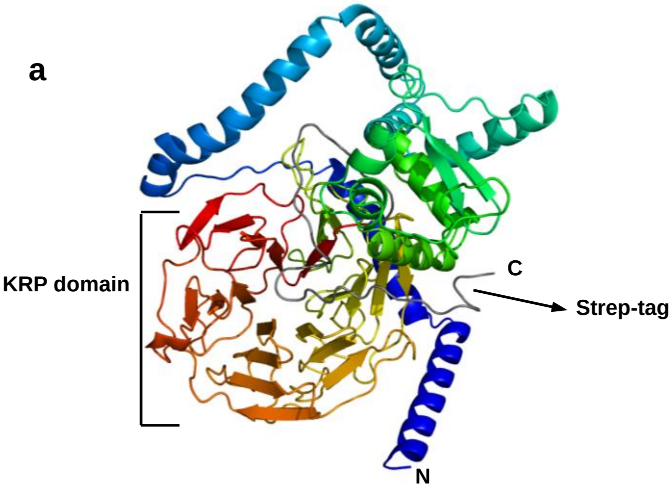


**b**

Truncating Trk13-WT-strep (572 amino acids) into 6 parts

```
KFSTVNNVNDTYEKKIIETELSDASDFENMVGDLRITFINWLKKTQMNFIREKDKLFKDKKELEMERVRLYKELENRKNIE
EQKLHDERKKLDIDISNGYKQIKKEKEEHRKRFDEERLRFLQEIDKIKLVLYLEKEKYYQEYKNFENDKKKIVDANIATET
MIDINVGGAIFETSRHTLTQQKDSFIEKLLSGRHHVTRDKQGRIFLDRDSELFRIILNFLRNPLTIPIPKDLSESEALLKE
AEFYGIKFLPFPLVFCIGGFDGVEYLNSMELLDISQQCWRMCTPMSTKKAYFGSAVLNNFLYVFGGNNYDYKALFETEVYD
RLRDVWYVSSNLNIPRRNNCGVTSNGRIYCIGGYDGSSIIPNVEAYDHRMKAWVEVAPLNTPRSSAMCVAFDNKIYVIGGT
NGERLNSIEVYEEKMNKWEQFPYALLEARSSGAAFNYLNQIYVVGGIDNEHNILDSVEQYQPFNKRWQFLNGVPEKKMNFG
AATLSDSYIITGGENGEVLNSCHFFSPDTNEWQLGPSLLVPRFGHSVLIANIAAGLSAWSHPQFEKGGGSGGGSGGGSWSH
PQFEK
```

1-23: Disordered region (Yellow)
24-42: Intervening region (Moss green)
53-99: Coiled-coil region 1 (Purple)
100-161: Coiled-coil region 2 (Red)
162-260: BTB/POZ region (Cyan)
261-572: Propeller region plus strep tag (Dark grey). The one-Strep tag sequence is underlined

**c**

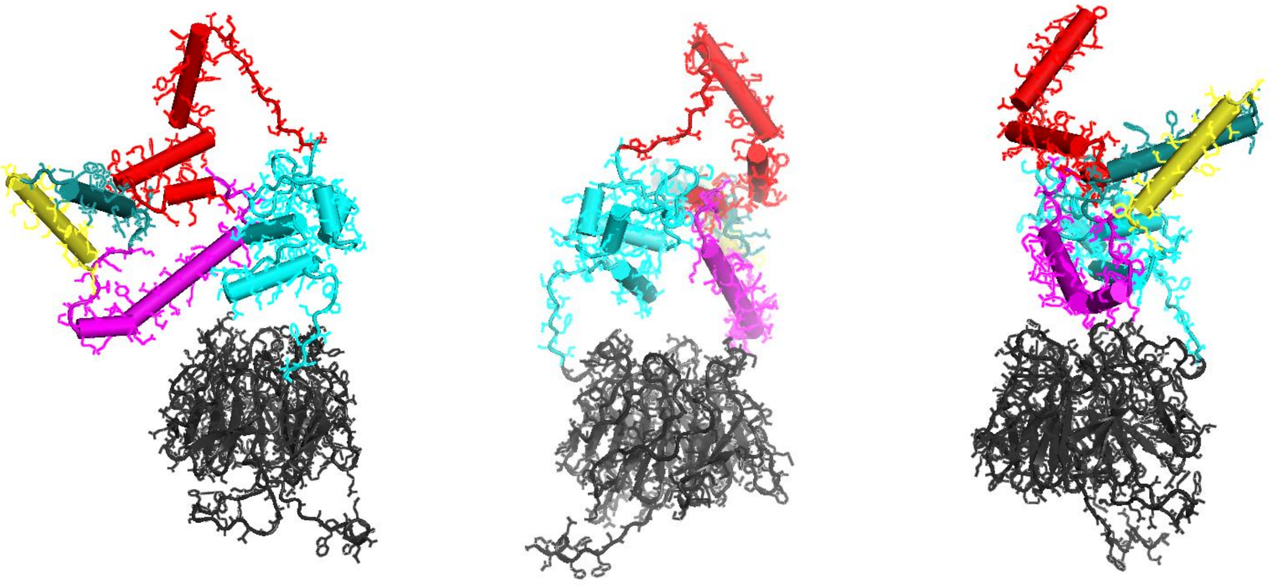

### Supplementary Figure 5

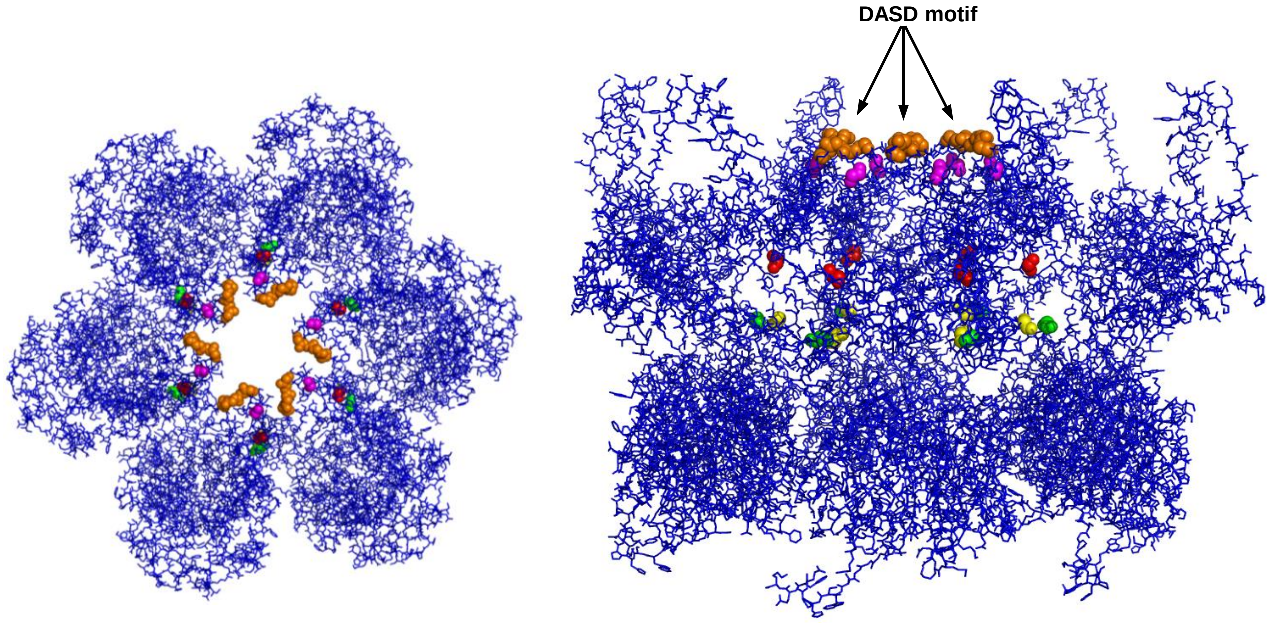
